# Oncogenic and immunomodulatory functions of SUV420H1 in HPV-negative head and neck squamous cell carcinoma

**DOI:** 10.1101/2025.10.20.683419

**Authors:** Arfa Moshiri, Marie Luff, Sohyoung Kim, Meiye Jiang, Mohd Saleem Dar, Malhar Patel, Katherine McKinnon, Elijah F. Edmondson, Jawad Akhtar, Hui Cheng, Vassiliki Saloura

## Abstract

Despite the advent of immunotherapy, human-papilloma-virus (HPV)-negative head and neck squamous cell carcinoma (HNSCC) carries a high morbidity and mortality rate, thus novel therapies are urgently needed. Suppressor Of Variegation 4-20 Homolog 1 (SUV420H1) is a protein lysine methyltransferase that writes H4K20me3. Approximately 35% of HPV-negative HNSCC tumors carry gains/amplifications of *SUV420H1*. Gene Set Enrichment Analysis (GSEA) showed enrichment of proliferation-, epithelial-mesenchymal transition (EMT)- and immune-response pathways in *SUV420H1*-overexpressing HPV-negative HNSCC tumors. Depletion of SUV420H1 led to decreased proliferation, cell cycling and invasion in human HPV-negative HNSCC cell lines, while enzymatic inhibition decreased the invasive capacity but not the proliferation and cell cycling of HPV-negative HNSCC cell lines, supporting the presence of catalytically-independent and -dependent functions of SUV420H1. In a syngeneic mouse model of mouse oral carcinoma 1 tumors (MOC1), *Suv420h1* knockout (KO) in MOC1 cancer cells halted tumor growth and synergized with anti-PD-1 therapy. In the tumor immune microenvironment (TIME) of *Suv420h1* KO MOC1 tumors, a significant increase in the macrophage compartment with a concurrent decrease in the protumorigenic granulocytic myeloid derived suppressor cells (gMDSCs) was observed. Genome-wide mapping of SUV420H1-mediated H4K20me3 revealed enrichment of EMT-, IFN-response, and myeloid-attracting chemokines. This work provides rationale for the depletion/degradation of SUV420H1 as a novel therapeutic strategy to hinder proliferation and invasion, and to impart sensitization to anti-PD-1 immunotherapy for patients with HPV-negative HNSCC tumors with *SUV420H1* gain/amplification.

## INTRODUCTION

Head and neck squamous cell carcinoma (HNSCC) represents a significant global health burden, encompassing a diverse group of malignancies originating from the mucosal linings of the oral cavity, pharynx, and larynx (1). There are two main subtypes of HNSCC, human-papilloma-virus (HPV)-positive and HPV-negative HNSCC; the latter has a significantly worse prognosis with recurrence rates of approximately 50% after standard of care surgery and/or chemoradiotherapy (2). Furthermore, despite recent advances in the treatment of recurrent/metastatic (R/M) HNSCC with the approval of pembrolizumab, an anti-programmed-death-1 antibody (anti-PD-1), the prognosis of patients with R/M HNSCC is still tenable, with a median overall survival of 13 months (3). These suboptimal outcomes emphasize the urgent need for more effective therapeutic strategies, particularly for HPV-negative HNSCC.

The Suppressor Of Variegation 4-20 Homolog 1 (SUV420H1 or KMT5B) is a key regulator of DNA replication and repair, chromatin structure and gene expression. It utilizes mono-methylated H4 lysine 20 (H4K20me1) as a substrate to catalyze its di- and tri-methylation (H4K20me2/3) (4), a modification intricately involved in genome integrity and DNA damage response, chromatin compaction and transcriptional regulation (5–12). Specifically, SUV420H1 has been found to facilitate nonhomologous end-joining (NHEJ)-directed DNA repair by catalyzing di- and tri-methylation of H4K20, promoting genomic stability (7, 9). Furthermore, Gonzalo et al showed that SUV420H1 cooperates with Retinoblastoma RB1 to maintain constitutive heterochromatin and the overall chromatin structure through H4K20me3 (8). The same group also demonstrated a role for SUV420H1 in the stability and integrity of telomeres through H4K20me3, showing that depletion of SUV420H1 induced telomere elongation and increased the frequency of telomere recombination (5). Interestingly, a recent study showed that SUV420H1 possesses both catalytic and non-catalytic cellular functions, and that its catalytic activity is enhanced by the histone variant H2A.Z (4). A non-catalytic function constitutes the binding of SUV420H1 on the chromatin, inducing a dramatic detachment of the DNA from the histone octamer, increasing nucleosomal breathing and chromatin compaction.

More recently, studies have investigated the role of SUV420H1 in cancer. Yokohama et al showed that SUV420H1-mediated H4K20me3 was associated with a better prognosis in breast cancer patients and that ectopic expression of SUV420H1 suppressed the invasive capacity of breast cancer cell lines, suggesting a tumor suppressor function in breast cancer (12). Accordingly, Lopez et al. demonstrated that ectopic overexpression of SUV420H1 induced tumor-suppressor-like features in vitro and that loss of SUV420H1 expression contributed to malignant transformation in glioblastoma multiforme (10). Furthermore, Brohm et al studied recurrent cancer mutations in SUV420H1 and found that 7 out of 8 of them induced a reduction in its catalytic activity, suggesting a tumor suppressive function (6). In contrast, other studies support the function of SUV420H1 as an oncogene in other cancer types, such as HPV-negative HNSCC and hepatocellular carcinoma (HCC) (11, 13). Specifically, Vougiouklakis et al showed that SUV420H1 depletion decreased the proliferation of HPV-negative HNSCC cell lines and that this was mediated both through ERK1 protein methylation and transcriptional regulation of the *ERK1* gene (11). Kato et al demonstrated that SUV420H1 is associated with poor prognosis in HCC and that its depletion reduced the proliferative and invasive capacity of HCC cell lines (13). The above suggest that the role of SUV420H1 in cancer is context-specific and that further investigation is needed to fully understand the mechanisms through which SUV420H1 mediates its tumor suppressive or oncogenic effects.

The present study aimed to investigate the significance of SUV420H1 as a novel therapeutic target in HPV-negative HNSCC. Through genetic depletion using siRNAs and clustered regularly interspaced short palindromic repeats (CRISPR) editing, pharmacological inhibition, and high-throughput genome-wide mapping of H4K20me3 through cleavage under targets and release using nuclease (CUT&RUN) assays, we unveil the role of SUV420H1 as an oncogene and immunomodulator in HPV-negative HNSCC. We identify distinct non-catalytically and catalytically-dependent phenotypes associated with SUV420H1 and putative direct gene targets regulated by SUV420H1-mediated H4K20me3 in HPV-negative HNSCC. This study provides insights into the role of SUV420H1 in HPV-negative HNSCC and underscores the rationale for SUV420H1-targeted therapeutic intervention for patients with SUV420H1-driven HPV-negative HNSCC.

## MATERIALS AND METHODS

### Generation of Suv420h1 knockout and HA-tagged catalytically-inactive SUV420H1 cell lines using CRISPR

*Suv420h1* knockout (KO) mouse oral carcinoma 1 (MOC1) cell lines (M02, M06) were generated from parental mouse MOC1 cells using clustered regularly interspaced short palindromic repeats (CRISPR/Cas9) technology, as previously described(20). H4k20me3 protein levels were assessed by Western blotting to confirm the efficiency of *Suv420h1* KO (**Fig. 6A**). HA-tagged catalytically-inactive *SUV420H1* cell lines were generated from human HN-6 cells using CRISPR/Cas9. Specifically, point mutations substituting histidine at 273aa position for lysine (H273K) as Mut1 or cysteine at 275aa position for arginine (C275A) as Mut2 were introduced in the wild-type expressing *SUV420H1* gene using HN-6 parental cells (diploid state for *SUV420H1*). Catalytic inactivity was confirmed by Western blotting for H4K20me3 levels (**Fig. 2H**).

### Cell culture

HPV-negative squamous cell carcinoma human cell lines HN-6, HN13, PE/CA-PJ15, HN-SCC-151 and FaDu cells were derived from patients with locoregionally advanced HPV-negative HNSCC and were kindly provided by Dr. Tanguy Seiwert (University of Chicago). HN-6 and HN13 cells were maintained in DMEM medium with 10% fetal bovine serum, 1% penicillin/streptomycin, and 2 nM L-glutamine. PE/CA-PJ15 cells were maintained IDMEM medium with 10% fetal bovine serum, 1% penicillin/streptomycin, and 2 nM L-glutamine. HN-SCC-151 cells were maintained in DMEM/F12 medium, 10% fetal bovine serum, 1% penicillin/streptomycin and 2 nM L-glutamine. FaDu cells were maintained in RPMI medium with 10% fetal bovine serum, 1% penicillin/streptomycin, and 2 nM L-glutamine.

MOC1 is a transplantable mouse HPV-negative squamous cell carcinoma cell line derived from carcinogen-induced primary tumors in C57BL/6 WT mice (15). MOC1 (parental cells), NT (sgRNA control), M02 and M06 (Suv420h1 KO) cell lines were maintained in a solution of 62% IMDM medium, 31% Ham’s F-12 nutrient mix, 5% fetal bovine serum, 1% penicillin/streptomycin, 40 ug/L hydrocortisone, 5 ug/L exponential growth factor (EGF), and 5 mg/L insulin.

### siRNA transfections

siRNA oligonucleotides were purchased from Millipore-Sigma (Burlington, MA) to target the human *SUV420H1* mRNA (SASI_Hs02_00348728, termed siSUV420H1#1, SASI_Hs01_00146572, termed siSUV420H1#2, SASI_Hs01_00146575, termed siSUV420H1#3). The negative control siRNA was purchased from Dharmacon (siRNA negative control Dharmacon ON-TARGET plus control pool, #D-001810-10-50, Horizon Discovery). Depending on the assay (CCK8, CFAs, cell cycle flow cytometry), HNSCC cells were plated overnight in 24-well-, 6-well-plates or 10cm dishes and were transfected with siRNA duplexes (50 nM final concentration) using Lipofectamine RNAimax (Thermo Fisher Scientific), with re-transfection performed every fourth day.

### CCK8 assays

HNSCC cells were plated overnight in 24-well plates (2-4 × 104 cells/well) and on the next day, they were transfected with SUV420H1 (siSUV420H1#1, siSUV420H1#2, siSUV420H1#3) or control siRNAs (50 nM final concentration) using Lipofectamine RNAimax (Thermo Fisher Scientific) for 9-12 days, with re-transfection performed on day 4. The number of viable cells was measured using the Cell Counting Kit-8 (Dojindo).

### Colony formation assays (CFAs)

HNSCC cells were plated overnight in 6-well plates (500 cells/well) and on the next day, they were transfected with SUV420H1 (siSUV420H1#1, siSUV420H1#2, siSUV420H1#3) or control siRNAs (50 nM final concentration) using Lipofectamine RNAimax (Thermo Fisher Scientific) for 8-12 days, with re-transfection performed on day 4. Once colonies were visible, they were stained with 0.01% (w/v) crystal violet (Sigma-Adrich, cat # HT901-8FOZ), washed with dH20 to remove excess stain, and left to dry. Colonies were counted using the ImageJ software (version 1.53k).

### Cell cycle analysis with BrdU

Cell cycle analysis was performed using the 5-bromo-20-deoxyuridine (BrdU) flow kit (BD Pharmingen™ FITC BrdU Flow Kit, cat#51-2354AK, BD Biosciences) according to the manufacturer’s instructions. Briefly, cells were seeded overnight in 10cm tissue culture dishes and treated with si*SUV420H1#2* and *#3* (50 nM) versus negative control siRNAs (as per above) using medium with 10% FBS for 72h. 2h before cell collection, cells were incubated with 10 μM BrdU. Cells were trypsinized, washed, fixed and permealized. Then cells were incubated with DNAase for 1 h at 37°C, and FITC-conjugated anti-BrdU antibody (dilution: 1:50) was added for 20 min at room temperature. Total DNA was stained with 7-amino-actinomycin D (7-AAD), followed by flow cytometric analysis. Cell cycle analysis was performed in HN-SCC-151 cells and HN13 cells. The same assay was performed in HN-SCC-151 cells treated with the SUV420H1 inhibitor A-196 (2.5uM) as described in the methods section titled “A-196 treatments”.

### Transwell invasion assays

For the invasion assay, 50uL of Matrigel (Corning™, cat # 354234) were placed on top of the transwell inserts with 8μm pores (Millicell®, cat #PI8P01250) and the transwell was placed in a 24-well plate and was solidified in a 37 °C incubator for 1-2 hours to form a thin gel layer before seeding 80,000 cells/200uL . HN-SCC-151 and HN13 cells treated with siNC or si*SUV420H1#2* and *#3*, or with DMSO or A-196 for 6 days, were seeded in the upper chambers (80,000 cells/200uL FBS free DMEM), and the lower chambers were filled with 600uL of FBS-containing DMEM culture medium. After 24 hours of incubation at 37°C, the upper side of the transwells was gently swabbed with a cotton swab to remove non-invaded cells, while the cells that had invaded through the Matrigel and transwell pores to the lower surface were fixed and stained with 500uL of 0.2% crystal violet for 1-2 hour at room temperature. The crystal violet was removed with a cotton swab and by dipping the transwells in dH20. The membranes were left to dry and migrated cells were imaged and counted using ImageJ (version 1.53k), Invasion assay was performed in biological triplicates and replicated once.

### A-196 treatments

To inhibit the enzymatic activity of SUV420H1, cells were treated with the A-196 inhibitor (MedChemExpress, Cat. No.: HY-100201). DMSO (Sigma-Aldrich, CAS No.: 67-68-5) was used as the vehicle control. Cells were plated in 24-well plates, 6-well plates, or 10 cm dishes depending on the assay, and treated with A-196 at final concentrations of 0.5 μM, 1 μM, or 2.5 μM. An equivalent volume of DMSO corresponding to that used for the highest A-196 concentration (2.5 μM) was applied to all control samples. Re-treatment with A-196 or DMSO was performed every 3 to 4 days.

### Western blotting

Nuclear extracts were prepared from the presented conditions using the Nuclear Complex Co-IP kit (Active Motif) and 5 μg were loaded to examine protein levels of H4K20me3, with histone H3 used as a loading control.

Primary antibodies used were anti-H4K20me3 (CST, 5737S, dilution 1:1000) and anti-histone 3 (ab1791, Abcam, Cambridge, MA, dilution 1:125,000). Amersham ECL prime Western Blotting Detection Reagent (Cytiva, cat # 45-002-401) or Pierce ECL Western Blotting Substrate (Thermo Scientific, cat# 32106) were used as detection reagents. Blots were imaged using a Chemi-fluorescent Odyssey FC machine after applying a detection reagent. Densitometry of all blots was performed using ImageJ software (1.53k, NIH, Bethesda, MD).

### Immunohistochemistry of HPV-negative tumors and normal buccal mucosal epithelium samples

As previously described (39), formalin-fixed, paraffin embedded tissue microarrays containing clinically annotated, de-identified patient tumor samples were obtained from the Human Tissue Research Center of the University of Chicago Pathology Department (IRB#8980). The IHC staining was approved by the Institutional Review Board of the University of Chicago (IRB#18-0468-AM002). 96 HPV-negative HSNCC tumors were stained for SUV420H1 (Abcam, cat#ab118659) using immunohistochemistry (IHC). The staining was performed on Leica Bond RX automated stainer. After deparaffinization and rehydration, tissue sections were treated with antigen retrieval solution (Leica Microsystems) with heat near 100°C for 20 minutes. The anti-SUV420H1 antibody (1:300) was applied on tissue sections for 1 hour incubation at room temperature. The antigen-antibody binding was detected with Leica Bond Polymer Refine Detection system (Leica Microsystems) and the slides were covered with cover glasses.

Expression of SUV420H1 was quantified using brightfield immunohistochemistry in tissue microarray slides containing normal and neoplastic squamous epithelium. Slides were digitalized at 20× objective (0.5 × 0.5µm per pixel) using Aperio AT2 scanner (Leica Biosystems) and analyzed using QuPath (v0.5.1). Appropriateness of staining and regions of interest annotation were completed by a board-certified pathologist to include viable tumor and exclude tumor necrosis, non-tumor tissues, and histology artifact. Quantification of tumor positivity was performed using image analysis, which included identifying stain vectors, optimizing cell detection algorithms, and thresholding chromogenic IHC. SUV420H1 staining was quantified separately for nuclear and cytoplasmic positivity; cells were binned based on staining intensity as negative, 1+, 2+, or 3+ and an H-score was calculated (Hirsch 2003).

### Quantitative real-time PCR

For quantitative real-time PCR, primers for *GAPDH* (housekeeping gene) and *SUV420H1* were purchased from Sigma-Aldrich. RNA extraction was performed using the RNeasy Mini Kit (cat # 74004, Qiagen Sciences Inc). cDNA conversion was performed using the Invitrogen SuperScript III First-Strand Synthesis System for RT-PCR (cat# 18080051, Invitrogen). PCR was conducted in technical triplicates using SYBR Select Master Mix (cat# 4472908, Applied Biosystems). PCR reactions were performed using the Applied Biosystems ViiA 7 machine (Thermo Fisher Scientific, Waltham, MA) and the QuantStudio Real-Time PCR Software v1.6.1 (Thermo Fisher Scientific). Subsequent analysis of the results was conducted using Microsoft Excel. and relative gene expression levels were calculated using the ΔΔCt method with GAPDH as the internal control.

### RNA-sequencing

RNA-seq was performed in HN-SCC-151 cells treated with siNC or si*SUV420H1#2* and *#3*, and DMSO or A-196 for 6 days. Specifically, cells were plated, treated with siRNAs or A-196 for 6 days and then they were trypsinized, washed twice with PBS, centrifuged and processed for RNA extraction (Direct-zol RNA miniprep kit, Zymo Research). Three biological replicates for each sample were processed to extract RNA and were quantified using Qubit. The Illumina® Stranded mRNA Prep kit was used per manufacturer’s protocol for library construction. Samples were pooled and sequenced on NextSeq 2000 P2 using Illumina® Stranded mRNA Prep and paired-end sequencing. The samples had 147-197 million pass filter reads. Reads of the samples were trimmed for adapters and low-quality bases using Cutadapt before alignment with the reference genome (hg38) and the annotated transcripts using STAR/RSEM. The samples had 59-61% non-duplicate reads.

### CUT&RUN assays and DNA-sequencing

For CUT&RUN assays, the 14-1048 CUT&RUN kit by EpiCypher was utilized according to EpiCypher’s protocol. Briefly, CUTANA spike-in dNuc controls (H3K4me0, 1, 2, 3) were mixed together with washed streptavidin (SA) beads in 4 separate 1.5ml tubes, and incubated for 30min at RT on nutator. Concanavalin (ConA) beads were activated using cold bead activation buffer, washed twice using a magnet, resuspended in cold activation buffer, added at 10uL/sample in separate strip tubes (1 tube per experimental sample) and kept on ice. 500,000 cells per experimental condition were obtained after trypsinization from respective cell culture dishes (1 dish per biological replicate) and were washed with PBS x 3 to remove excess trypsin. Cells were then resuspended in 100uL/sample of RT wash buffer and washed twice at 600xg for 3min. After the final wash, cell pellets were resuspended in 105uL of RT wash buffer, and 100uL per sample were aliquoted into each 8-strip tube containing 10uL of activated beads. The cell-bead slurries were incubated on the benchtop for 10min at RT to allow for adsorption of the cells to the beads. After the incubation, the slurries were placed on a magnet and a small aliquot of the supernatant was obtained to confirm adsorption of cells to the beads (binding efficacy >93% of cell input). The supernatants were completed removed and the cell-bead slurries were then exposed to cold antibody buffer and vortexed. The CUTANA H3K4MetSTat spike-in control dNucs were added to designated positive (H3K4me3) and negative (IgG) control tubes. Then, 0.5ug of antibody targeting H4K20me3 (CST, 5737S) was added to each designated experimental tube. Biological triplicates were used for each experimental condition. The samples were incubated overnight on a nutator at 4oC. Next day, the 8-strip tubes containing the samples were placed on a magnet till the slurries cleared, supernatants were removed and the cell-beads were washed twice with cold cell permealization buffer. After the final wash, 50uL of the cold cell permealization buffer was added to the cell-bead slurries, and then 2.5uL of pAG-Mnase was added to each sample. Samples were incubated for 10min at RT and the 8-strip tubes were placed back on a magnet. Supernatants were removed and cell-beads complexes were washed twice with cold cell permealization buffer. After the final wash, 50uL of cell cell permealization buffer was added in each sample, and targeted chromatin digestion followed by adding 1uL of chromatin digest additive to each sample. Strips were incubated for 2h at 4oC on a nutator and the reaction was stopped using Stop buffer. 0.5ng of spike-in Ecoli DNA was added to each sample and samples were incubated for 10min at 37oC in a thermocycler. The strips were then placed on a magnet and the supernatants containing the CUT&RUN enriched DNA were transferred to new tubes. DNA was purified per EpiCypher’s protocol, and library construction was conducted. Nucleic acid size selection to enrich for fragment sizes between 200-500bp was conducted using SPRIselect (Beckman Coulter Life Sciences, B23318). Samples were pooled and sequenced on NextSeq2000 using TruSeq ChIP and Swift Bioscience Accel-NGS 2S Plus DNA Library Prep Kits and paired-end sequencing. All the samples had yields between 54 and 87 million pass filter reads. Samples were trimmed for adapters using Cutadapt before the alignment. The trimmed reads were aligned with hg38 reference using Bowtie2 alignment. All the samples had library complexity with percent non-duplicated reads ranging from 68 to 76%.

### Mouse experiments

All animal experimental protocols, study designs and animal usage were approved and conducted accordingly to all applicable guidelines by the NCI-Bethesda Animal Care and Use Committee and adhering to ARRIVE guidelines. 6-6 week-old female C57BL/6 mice were purchased from Taconic and used for the described experiments. Control NT MOC1, and *Suv420h1* KO cell lines M02 (KO-1) and M06 (KO-2) in vitro and were inoculated by subcutaneous injections of 5 million cells in suspension using Matrigel, in the right flanks of C57BL/6 mice. 6-7 C57BL/6 mice were injected per group. Tumor length (L) and width (W) were measured twice weekly with calipers starting from the day of inoculation and tumor volumes were calculated using the formula LxW^2/2. 30 days following tumor inoculation, the mice were euthanized using cervical dislocation, tumors were surgically resected, processed to single-cell suspensions (Miltenyi Biotec) and processed for flow cytometry as described below.

### Multicolor flow cytometry

For the multicolor flow of NT, KO-1 and KO-2 MOC1 tumors, mice were euthanized and flank MOC1 tumors were surgically resected, and mechanically and chemically digested into single-cell suspensions using the gentleMACS Dissociator and the mouse tumor dissociation kit (Miltenyi Biotec), per manufacturer’s protocol. Single-cell suspensions were filtered through 70um filters and washed with 1% BSA in PBS. Samples were incubated with anti-CD16/32 (Biolegend) antibody to block nonspecific staining. Subsequently, the primary antibodies were added and incubation for 30min was carried out in the dark. Cell surface staining was performed using fluorophore-conjugated Alexa 488 anti-mouse PDL1 (BD 566864), PE-Dazzle 594 PD1 (BioLegend 135228), PE-Cy5 PDGFR/CD140a (BioLegend 135920), PE-Cy7 I-Ab/MHC II (BioLegend 116420), APC CD11b (BioLegend 101212), Alexa 700 CD3e (BioLegend 152316), APC-Cy7 CD25 (BioLegend 102026), BV510 F4/80 (BioLegend 123135), BV605 CD11c (BioLegend117334), BV750 CD31 (BD 746871), BV786 CD45.2 (BioLegend 109839), BUV661 CD8a (BD 750023), BUV737 H2-Kb/MHC I (BD 748822), and BUV805 CD4 (BD 612900). Cell viability was assessed with LIVE/DEAD Fixable Blue Dead Cell Stain Kit (L23105, Invitrogen, Waltham, MA). All analyses was performed on a BD FACSymphony A5 analyzer (BD Biosceinces, Franklin Lakes, NJ) running FACSDiva software and interpreted using FlowJo V.X10.0.7r2.

### Bioinformatics and Statistical Analysis

#### GSEA analysis of TCGA datasets

For the gene-set enrichment analysis (GSEA), 422 TCGA HPV-negative tumor samples (Firehose Legacy) with mRNA expression data (data_mrna_seq_v2_rsem.txt) were analyzed and a ranked gene list was obtained based on Pearson’s correlation of each gene with *SUV420H1* expression (**Fig. 1C**). Using the R package gsea (v1.32.0), GSEA was performed on the ranked gene list, with MsigDB’s Hallmark gene sets from human (obtained via R package msigdb v1.14.0) serving as the reference.

**Figure 1.**
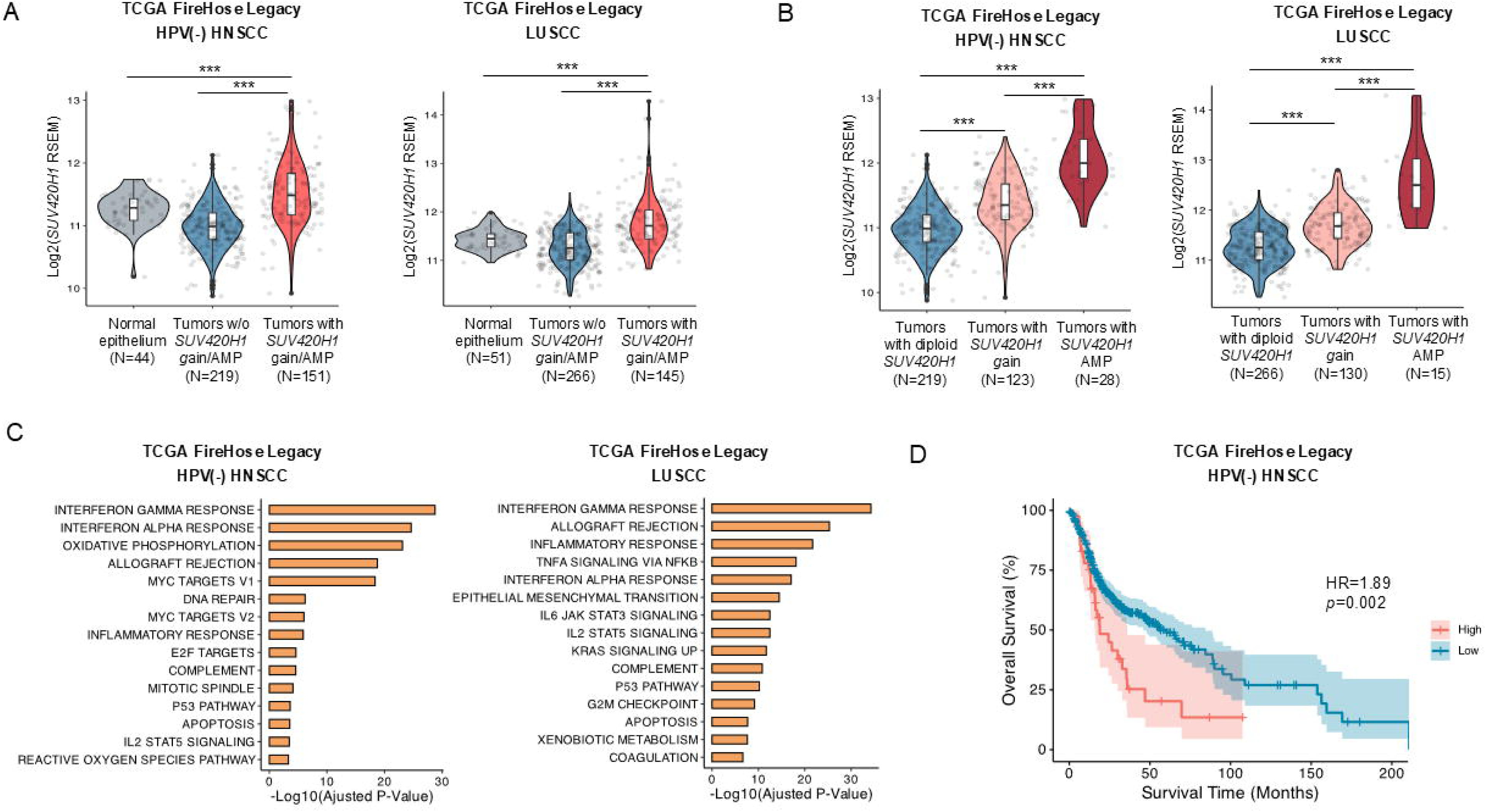
*SUV420H1* is amplified and is associated with worse overall survival in HPV-negative HNSCC. **(A)** Violin plots comparing *SUV420H1* mRNA expression between normal epithelium (buccal mucosa or bronchial) and HPV-negative HNSCC (left) or lung SCC (LUSCC) tumors (right) with or without *SUV420H1* gain/amplification. Firehose Legacy, TCGA. Wilcoxon rank-sum test; * p<0.05, ** p<0.01, *** p<0.001. **(B)** Correlation of copy number gain and amplification with mRNA expression of *SUV420H1* in HPV-negative HNSCC (left) and LUSCC (right). Firehose Legacy, TCGA. Wilcoxon rank-sum test, * p<0.05, ** p<0.01, *** p<0.001. **(C)** Gene Set Enrichment Analysis (GSEA) using the HPV-negative HNSCC (left) and LUSCC (right) cohorts of the Firehose Legacy, TCGA. *SUV420H1* mRNA expression is associated with IFN-related, cell cycle and epithelial-mesenchymal transition (EMT) gene signatures in patients with HPV-negative HSNCC and LUSCC. The GSEA was performed on a ranked gene list. Genes were ranked by their Pearson correlation with *SUV420H1* mRNA. The top 15 pathways are presented here based on adjusted p-values. **(D)** Kaplan-Meier overall survival curve of HPV-negative HNSCC patients dichotomized by *SUV420H1* mRNA expression levels (TCGA). Log-rank p-value. HR: hazard ratio. High group was defined by the top 10% of *SUV420H1* mRNA expression.

**Figure 2.**
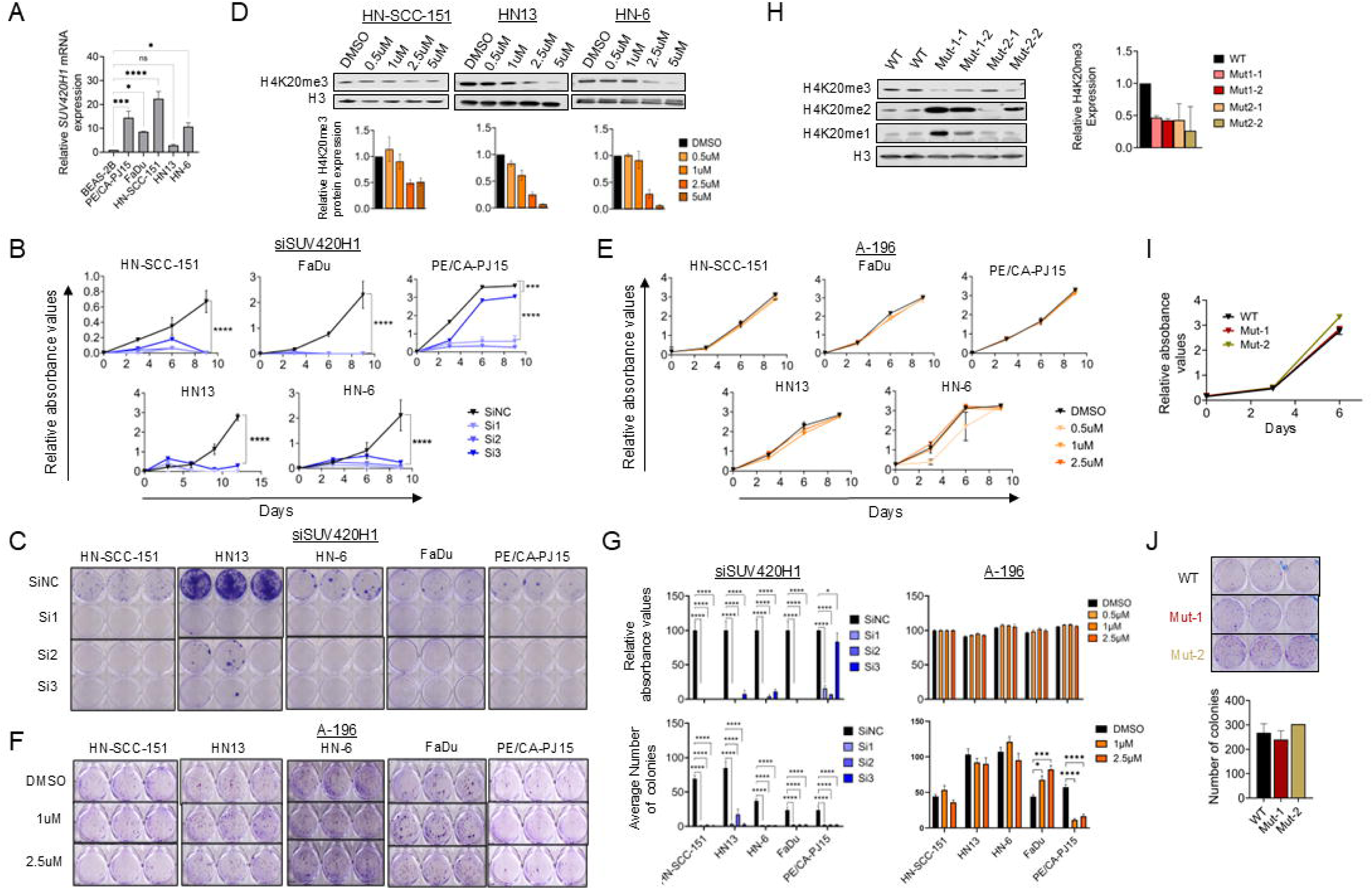
Effect of SUV420H1 depletion and enzymatic inhibition on the proliferative capacity of HPV-negative HNSCC cells. **(A)** Relative *SUV420H1* mRNA expression levels in the immortalized squamous epithelial cell line BEAS-2B and 5 HPV-negative HNSCC cell lines (PE/CA-PJ15, FaDu, HN-SCC-151, HN13, HN-6). Data are presented as mean ± SEM from n = 3 biological replicates. Ordinary one-way ANOVA with multiple comparisons; * p<0.05, ** p< 0.01, *** p<0.001, **** p<0.0001. **(B)** CCK8 assays in 5 HPV-negative HNSCC cell lines treated with negative control (siNC) or three *SUV420H1*-targeting siRNAs (si1, si2, si3) for 10-12 days. Data are presented as mean ± SEM from n=5 biological replicates. Student t-test; * p<0.05, ** p< 0.01, *** p<0.001, **** p<0.0001. **(C)** Colony forming assays (CFAs) in 5 HPV-negative HNSCC cell lines treated with negative control (siNC) or three *SUV420H1*-targeting siRNAs (si1, si2, si3) for 10 days. **(D)** Top: Western blotting for H4K20me3 levels in HN-SCC-151, HN13 and HN-6 cells treated with control DMSO or incremental concentrations of the SUV420H1 inhibitor A-196 for 6 days. Nuclear extracts were obtained and 5ug were loaded. H3 was used as a loading control. Bottom: Densitometry for H4K20me3 bands, normalized by H3 band intensity. Data are shown as mean ± SEM from n=2 independent biological replicates. **(E)** CCK8 assays in 5 HPV-negative HNSCC cell lines treated with control DMSO or A-196 at 0.5uM, 1uM and 2.5uM for 12 days. Data are presented as mean ± SEM from n=5 biological replicates. Student t-test; * p<0.05, ** p<0.01, *** p<0.005. **(F)** Colony forming assays (CFAs) in 5 HPV-negative HNSCC cell lines treated with control DMSO or A-196 at 0.5uM, 1uM and 2.5uM for 12 days. **(G)** Top: Bar graphs of quantified results of CCK8 assay after 10 days of treatment of 5 HPV-negative HNSCC cell lines with *SUV420H1*-targeting siRNAs (left) or with A-196 (right). Data are presented as mean ± SEM from n=5 biological replicates. Bottom: Bar graphs of quantified results of CFAs after 10 days of treatment of 5 HPV-negative HNSCC cell lines with *SUV420H1*-targeting siRNAs (left) or with A-196 (right). Data are presented as mean ± SEM from n=3 biological replicates. Student t-test; * p<0.05, ** p<0.01, *** p<0.001, **** p<0.0001. **(H)** Top: Western blotting for H4K20me3, H4K20me2 and H4K20me1 in wild-type HN-6 parental cells and *SUV420H1*-mutant cell lines Mut-1 (H273K) and Mut-2 (C275A). 5ug of nuclear extracts were loaded. H3 was used as a loading control. Two biological replicates are presented. Bottom: Densitometry for H4K20me3 bands, normalized by H3 band intensity. Data are shown as mean ± SEM from n=2 biological replicates. **(I)** CCK8 assay with parental wild-type HN-6 cells and *SUV420H1*-mutant cell lines Mut-1 and Mut-2. Data are shown as mean ± SEM from n=5 biological replicates. **(J)** Top: CFAs of wild-type HN-6 parental cells and *SUV420H1*-mutant cell lines Mut-1 and Mut-2. Bottom: Bar graphs of quantified results of CFAs. Data are presented as mean ± SEM from n=5 biological replicates.

#### Clinical correlations

Cox regression was performed to examine whether *SUV420H1* mRNA levels (TCGA, FireHose Legacy, HPV-HNSCC) were prognostic for overall survival (**Fig. 1D**). The *SUV420H1* mRNA expression data was dichotomized into high and low groups by systematically evaluating percentile cutoffs ranging from the top 10% to 90%. The threshold demonstrating the strongest statistical significance with overall survival (OS) outcomes was designated as the optimal high-expression group definition. Kaplan-Meier analysis was then performed to derive OS curves utilizing R packages survival (v3.8.3) and survminer (v0.5.0). The Cox proportional hazards model was used to estimate hazard ratios (HRs) with 95% confidence intervals (CIs).

#### GSEA analysis of RNA-seq datasets of HN-SCC-151 cells treated with siSUV420H1 or A-196 inhibitor

Top enriched pathways (adjusted p-value < 0.05) obtained in Gene Set Enrichment Analysis (GSEA) of SMYD3 KO 5-3 vs. HN-6 cell lines using fgsea (1.22.0) R library and hallmark gene sets (h.all.v7.4.symbols.gmt). GSEA output data were used for the barplot (**Fig. 5A**).

#### RNA-seq heatmaps and differentially expressed genes

RNA-seq data were quantitated to obtain raw tag counts of exon regions at the gene level using featureCount (**Fig. 5B-D**). The raw tag count data was variance stabilizing transformed using VST function in DESeq2 R library, and z-score of the transformed data was obtained to color code for heatmap. For clustered heatmaps, pheatmap R library was used with Euclidean distance and ward.D2 clustering options. Significance of gene expression changes was evaluated using DESeq2 R library and determined based on Wald-statistics (FDR<0.1) available from DESeq2 library for A-196 treated samples, and FDR<0.1 and log2 fold-change (>log2(1.3), <-log2(1.3)) for si*SUV420H1* treated samples.

#### CUT&RUN analysis

Raw fastq files were trimmed with Cutadapt (v 4.4) and aligned using bowtie (v 2-2.5.1) against hg38. For spike-in controls, the trimmed FASTQ files were aligned against the Escherichia coli MG1655 genome. Duplicated reads were removed using Picard tools (v 2.27.3), and normalization factors were derived based on the uniquely mapped fragments in the corresponding spike-in control data. The enriched regions with H4K20me3 signals were identified using GoPeaks with –broad option and FDR=0.05 options. Raw tag counts were obtained using bedtools (v 2.30.0) and the bedmap function in BEDOPS toolkit (v 2.4.41) for merged peak sites. The raw tag count data was normalized using the normalization factors derived from the spike-in control data, and then differential analysis was performed using DESeq2 to obtain a final list of regions for downstream analysis (FDR<0.1). No LFC threshold was applied for CUT&RUN data.

#### Annotation of peaks (CUT&RUN)

Each peak/enriched region in CUT&RUN assays was annotated with the nearby gene displaying the shortest distance between TSS and the center of each peak using the ChipSeeker (v 1.32.0, GENCODE V39).

#### Volcano plots

For all volcano plots, EnhancedVolcano (v 1.14.0) R library was used.

#### Statistical analyses of in vitro and in vivo assays results

Averages of at least three biological replicates from each control and experimental condition were obtained for MTT assays, CFAs, flow cytometry and invasion assays. For the in vivo mouse experiments, averages of tumor volumes were obtained from 6-7 mice per group. Student t-test was used to conduct statistical evaluations between control and experimental conditions.

## RESULTS

### *SUV420H1* is amplified and associated with worse overall survival in HPV-negative HNSCC patients

In a recently published study from our group, we identified protein methyltransferases/demethylases with recurrent genetic amplifications and thus with putative oncogenic function in HPV-negative HNSCC (14). *SUV420H1* was identified as the second most frequently recurrently amplified gene in this family of enzymes (after *NSD3*), with approximately 6% of HPV-negative HNSCC tumors having focal amplifications in the 11q13.2 chromosomal locus carrying *SUV420H1*. Importantly, when gains and amplifications of *SUV420H1* are considered together, the combined gain/amplification rate is ∼36% (**Supplementary Fig.1**). Amongst other squamous cell carcinomas (SCCs) which have a similar genetic background as HPV-negative HNSCC (15), a similar gain/amplification rate of *SUV420H1* was also observed in lung (∼28%) and bladder urothelial carcinoma (∼27%), with a lower rate in cervical SCC (12%) (**Supplementary Fig. 1**).

The presence of *SUV420H1* gain/amplification in HPV-negative HNSCC tumors was associated with significantly higher mRNA expression of *SUV420H1* compared to its expression in normal buccal epithelium (**Fig. 1A**). HPV-negative HNSCC tumors without *SUV420H1* gain/amplification did not demonstrate increased *SUV420H1* mRNA expression (**Fig. 1A**), supporting that the gain/amplification of *SUV420H1* is the main mechanism driving the overexpression of *SUV420H1*. This finding was further validated in the lung SCC cohort of the TCGA (**Fig. 1A**). Accordingly, *SUV420H1* mRNA expression was higher in HPV-negative HNSCC and lung SCC tumors with *SUV420H1* gain, and even higher in tumors with *SUV420H1* amplification compared to tumors that were diploid for the *SUV420H1* gene (**Fig. 1B**).

To determine the subcellular localization of SUV420H1 in HPV-negative HNSCC, immunohistochemical analysis of SUV420H1 was conducted using tissue microarrays of HPV-negative HNSCC tumors (n=96, University of Chicago). Results showed that although cytoplasmic localization was sometimes observed, SUV420H1 had a predominantly nuclear subcellular localization in cancer cells, suggesting a predominantly nuclear function (**Supplementary Fig. 2A, 2B, Supplementary Table 1**). Furthermore, SUV420H1 protein levels were significantly higher in the tumor compared to the stroma compartment of HPV-negative HNSCC tissue sections (**Supplementary Fig. 2C, Supplementary Table 1**).

To assess the pathways associated with *SUV420H1* mRNA expression levels in HPV-negative HNSCC tumors, Gene Set Enrichment Analysis (GSEA) was conducted using the HPV-negative HNSCC cohort of the TCGA (Firehose Legacy). Results showed enrichment in interferon-alpha (IFNA) and gamma (IFNG) response, cell cycle and epithelial-mesenchymal transition (EMT) pathways (**Fig. 1C, Supplementary Fig. 3**). Similar pathways were enriched in lung SCC tumors (TCGA, Firehose Legacy) (**Fig. 1C**). Finally, Kaplan-Meier overall survival (OS) analysis revealed that higher *SUV420H1* mRNA levels were associated with worse OS in HPV-negative HNSCC patients (**Fig. 1D**). The above data suggest a potential role of SUV420H1 as an oncogene in the subset of HPV-negative HNSCC patients with *SUV420H1* gain/amplification.

### SUV420H1 depletion reduces the proliferative and colony forming capacity of HPV-negative HNSCC cells in a catalytically-independent manner

To assess the effect of SUV420H1 on the proliferative capacity of HPV-negative HNSCC cells, CCK8 and colony forming assays (CFAs) were pursued in five HPV-negative HNSCC cell lines with endogenous mRNA overexpression of *SUV420H1* (HN-6, HN13, FaDu, PE/CA-PJ15, HN-SCC-151) compared to a control normalized bronchial epithelial cell line (BEAS-2B) (**Fig. 2A**). To assess whether depletion of SUV420H1 affects the proliferation of HPV-negative HNSCC cells, we utilized three different siRNAs targeting SUV420H1, but not other H4K20 methyltransferases, such as SUV420H2 and SETD8 (**Supplementary Fig. 4**). SiRNA-mediated SUV420H1 depletion induced a significant and marked decrease in the relative cell numbers (**Fig. 2B, 2G**) and colony forming capacity (**Fig. 2C, 2G**) by approximately 80-100% after 9-12 days of treatment in all five HPV-negative HNSCC cell lines.

To evaluate the importance of the catalytic activity of SUV420H1 in the proliferation of HPV-negative HNSCC cells, CCK8 and CFA assays were conducted with the SUV420H1 enzymatic inhibitor A-196 in the same panel of cell lines. To determine whether A-196 inhibits SUV420H1 in HPV-negative HNSCC cells, three HPV-negative HNSCC cell lines were treated with different concentrations of A-196 (0-5uM) for 6 days and global levels of H4K20me3, the enzymatic end product of SUV420H1, were assessed by Western blotting. A dose-dependent reduction of global levels of H4K20me3 was observed with gradually increasing concentrations of A-196 in HN-SCC-151, HN13 and HN-6 cells, with a 50-80% reduction observed at 2.5uM in all three cell lines and up to nearly 90% at 5uM in HN13 and HN-6 cells (**Fig. 2D**). For the CCK8 and CFAs, cells were treated with a range of 0-2.5uM of A-196. The concentration of 5uM was not tested to avoid the possibility of off-target effects at this higher concentration which had a similar effect in the global H4K20me3 levels as the concentration of 2.5uM. Interestingly, in these cell lines, as well as in FaDu and PE/CA-PJ15 cells, no effect was observed in the relative cell absorbance at 2.5uM of A-196 (**Fig. 2E, 2G**). Accordingly, the colony forming capacity of all cell lines was largely unaffected (**Fig. 2F, 2G**), with the exception of PE/CA-PJ15 cells with a nearly 90% decrease in the colony forming capacity of these cells (**Fig. 2F, 2G**). With the exception of this cell line, these results, when compared to the effect of siRNA-mediated SUV420H1 depletion inducing ∼90-100% decrease in the proliferative and colony forming capacity, support that the catalytic activity of SUV420H1 is dispensable for the proliferation of the majority of HPV-negative HNSCC cell lines.

To further validate these findings, CRISPR knock-in HPV-negative HNSCC cell lines endogenously expressing two variants of HA-tagged catalytically-inactive SUV420H1 were generated. Specifically, point mutations substituting histidine at position 273 with lysine (H273K, Mut-1) or cysteine at position 275 with alanine (C275A, Mut-2) were introduced into the wild-type *SUV420H1* gene in HN-6 parental cells (diploid state for *SUV420H1*). Global H4K20me3 protein levels were markedly decreased by approximately 80% in the cell lines expressing mutant SUV420H1, validating the reduced catalytic activity of these SUV420H1 mutant variants (**Fig. 2H**). Consistent with the results with the A-196 SUV420H1 inhibitor, CCK8 assays and CFAs with the two catalytically-inactive *SUV420H1* mutant expressing cell lines showed no significant decrease in the proliferative and colony forming capacity of these cells (**Fig. 2I, 2J**).

These results support that SUV420H1 drives the proliferative capacity of HPV-negative HSNCC cells and that this effect is mediated through a catalytically-independent mechanism.

### SUV420H1 depletion but not enzymatic inhibition induces cell cycle arrest in HPV-negative HNSCC cells

To assess whether SUV420H1 affected the proliferative capacity of HPV-negative HNSCC cells through an effect on the cell cycle progression, cell cycle flow cytometry was conducted in HN-SCC-151 cells treated with control and two SUV420H1-targeting siRNAs for 72h. SUV420H1 depletion induced a significant decrease in the G0/G1- and S-phases and arrest in the G2/M phase (**Fig. 3A, B**). Interestingly, treatment of HN-SCC151 cells with the SUV420H1 inhibitor A-196 did not have any effect on the cell cycle progression. Similar results were reproduced in HN13 cells, as well as in the catalytically-inactive *SUV420H1* mutant cell lines (**Fig. 3A-C**). These findings suggest that the effect of SUV420H1 on the cell cycle is mediated through a catalytically-independent mechanism, as observed with its effect on the proliferative capacity of HPV-negative HNSCC cells (**Fig. 2B-J**).

**Figure 3.**
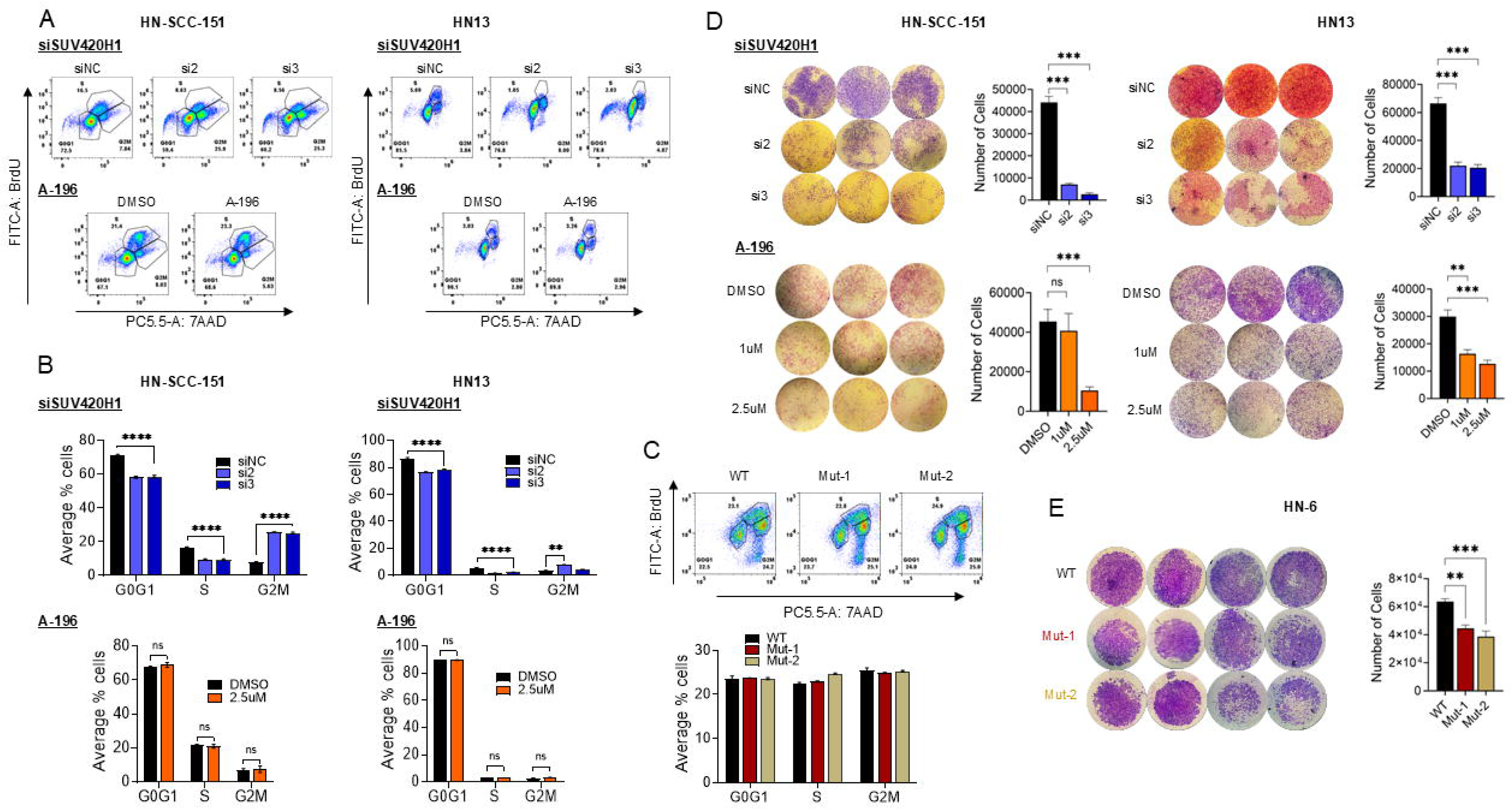
Effects of SUV420H1 depletion or enzymatic inhibition on the cell cycle progression and invasive capacity of HPV-negative HNSCC cells. **(A)** Cell cycle flow cytometry of HN-SCC-151 and HN13 cells treated with negative control (siNC) or *SUV420H1*-targeting siRNAs (si2, si3), and with DMSO or the SUV420H1 inhibitor A-196 at 2.5uM. Each graph is a representative example of three biological replicates. **(B)** Bar graph of quantified results of cell cycle flow cytometry of HN-SCC-151 and HN13 cells shown and described in (A). Data are presented as mean ± SEM from n=3 biological replicates. Student t-test; * p<0.05, ** p<0.01, *** p<0.001, **** p<0.001. **(C)** Bar graph of quantified results of cell cycle flow cytometry with catalytically inactive *SUV420H1*-mutant cell lines Mut-1 and Mut-2 compared to parental HN-6 cells. Data are presented as mean ± SEM from n=3 biological replicates. Student t-test; * p<0.05, ** p<0.01, *** p<0.001, **** p<0.001. **(D)** Invasion assays of HN-SCC-151 and HN13 cells treated with control (siNC) or *SUV420H1*-targeting siRNAs (si2, si3), and with DMSO or the SUV420H1 inhibitor A-196 at 2.5uM. Bar graph of quantified results of the invasion assay using Image J. Data are presented as mean ± SEM from n=3 biological replicates. Student t-test; * p<0.05, ** p<0.01,*** p<0.001. **(E)** Invasion assays of catalytically inactive *SUV420H1*-mutant cell lines Mut-1 and Mut-2 compared to parental HN-6 cells. Bar graph of quantified results of the invasion assay using Image J. Data are presented as mean ± SEM from n=3 biological replicates. Student t-test, * p<0.05, ** p<0.01, *** p<0.001.

### SUV420H1 depletion or enzymatic inhibition decreases the invasive potential of HPV-negative HNSCC cells

To evaluate the effect of SUV420H1 on the invasive potential of HPV-negative HNSCC cells, invasion assays were conducted using *SUV420H1*-targeting siRNAs, the SUV420H1 enzymatic inhibitor A-196 and the catalytically-inactive *SUV420H1* mutant cell lines. SiRNA-mediated depletion of *SUV420H1* induced a significant decrease in the invasiveness of HN-SCC-151 cells by approximately 90% (**Fig. 3D, top**). Interestingly, in contrast to the lack of efficacy seen in the proliferative potential and cell cycle progression of these cell lines, A-196 treatment induced a significant decrease in the invasiveness of these cells by approximately 80% at 2.5uM, similar to that observed with siRNA-mediated *SUV420H1* depletion (**Fig. 3D, bottom**). Similar results were obtained with HN13 cells (**Fig. 3D**). To further validate these results, invasion assays were also conducted with the catalytically-inactive *SUV420H1* mutant cell lines which accordingly showed a significant decrease in their invasive potential by ∼30-40% compared to the wild type parental cell line HN-6 (**Fig. 3E**). These results support that the invasive capacity of HPV-negative HNSCC cells is dependent on the catalytic activity of SUV420H1.

### *Suv420h1* knockout attenuates tumor growth, increases the macrophage intratumoral compartment and sensitizes MOC1 tumors to anti-PD-1 immunotherapy

Given the standard of care role of anti-PD-1 immunotherapy in the treatment of HPV-negative HNSCC (3) and the fact that IFN-response signatures were enriched in human tumors with *SUV420H1* mRNA overexpression (**Fig. 1C**), we opted to evaluate the effect of Suv420h1 in the tumor immune microenvironment (TIME) of mouse HPV-negative HNSCC tumors in vivo. To this purpose, *Suv420h1* knockout (KO) mouse oral carcinoma 1 (MOC1) cell lines were generated using CRISPR editing (**Fig. 4A**). The HPV-negative HNSCC MOC1 cell line has been previously established by exposure of the oral cavity of C57BL/6 mice to DMBA (7,12-dimethylbenz(a)anthracene) and is typically used to generate flank MOC1 tumors in a syngeneic, heterotopic C57BL/6 mouse model (16). In accordance with the results with human HPV-negative HNSCC cell lines (**Fig. 2B, 2C, 2G**), the colony forming capacity of two *Suv420h1* KO MOC1 cell lines (KO-1, KO-2) (**Fig. 4A**), as well as the global levels of H4k20me3 levels were decreased (**Fig. 4B**) compared to the parental MOC1 cells (negative control, NT).

**Figure 4.**
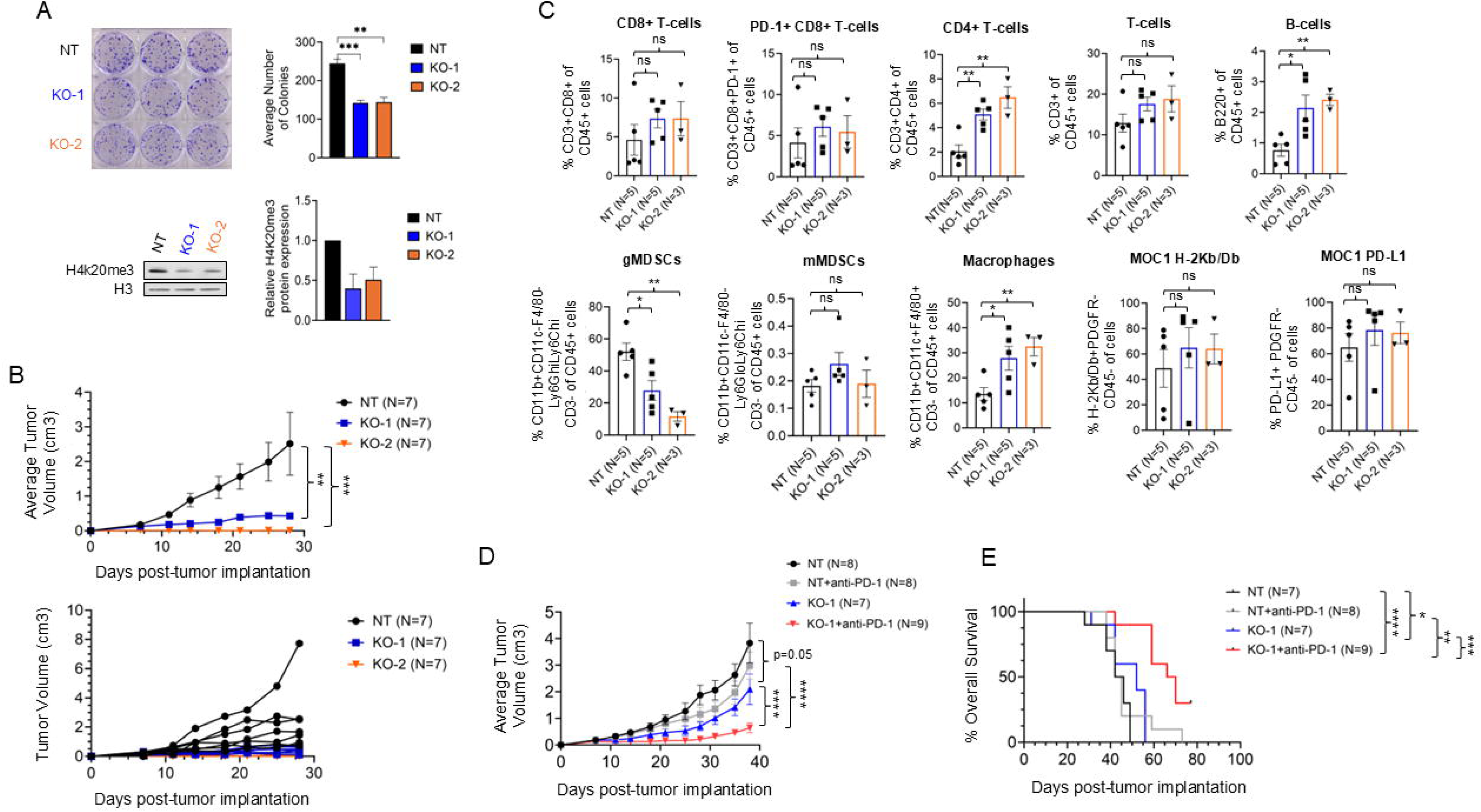
*Suv420h1* KO restraints tumor growth and sensitizes MOC1 tumors to anti-PD-1 therapy through macrophage influx. **(A)** *Suv420h1* knockout (KO) MOC1 cell lines generated with CRISPR. Top: Colony formation assays with control (NT) and two *Suv420h1* KO MOC1 cell lines (KO-1, KO-2). The graph represents the average number of colonies per condition. Data are shown as mean ± SEM from n=3 independent biological replicates. Bottom: Western blotting for H4k20me3 in control and *Suv420h1* KO MOC1 cell lines. Data are shown as mean ± SEM from n=2 biological replicates. **(B)** C57BL/6 mice were implanted with flank NT or *Suv420h1* KO-1 or KO-2 MOC1 tumors and tumor volumes were measured over 28 days. Top graph: Average tumor volume (cm3). Data are shown as mean ± SEM. Two-way ANOVA test; ** p<0.01, *** p<0.001. Bottom graph: hairline tumor growth curves. **(C)** Multicolor flow cytometry analysis of wild-type and *Suv420h1* KO MOC1 tumors. Data are represented as mean ± SEM. Unpaired t-test between NT and KO-1 or NT and KO-2; * p<0.05, ** p<0.01, *** p<0.001. **(D)** Average tumor volume (cm3) measured over 24 days in C57BL/6 mice bearing flank tumors from four experimental groups: control (NT), NT treated with anti-PD-1 antibody (NT+anti-PD-1), *Suv420h1* KO-1, and *Suv420h1* KO-1 treated with anti-PD-1 (KO-1+anti-PD-1). Two-way ANOVA test; p=0.05, **** p<0.0001. **(E)** Overall survival curves of four mouse groups: NT (n=7), NT+anti-PD-1 (n=8), *Suv420h1* KO-1 (n=7), and *Suv420h1* KO-1+anti-PD-1 (n=9). Survival was monitored for up to 80 days post-tumor implantation. Log-rank test; * p<0.05, ** p<0.01, *** p<0.001, **** p<0.0001.

**Figure 5.**
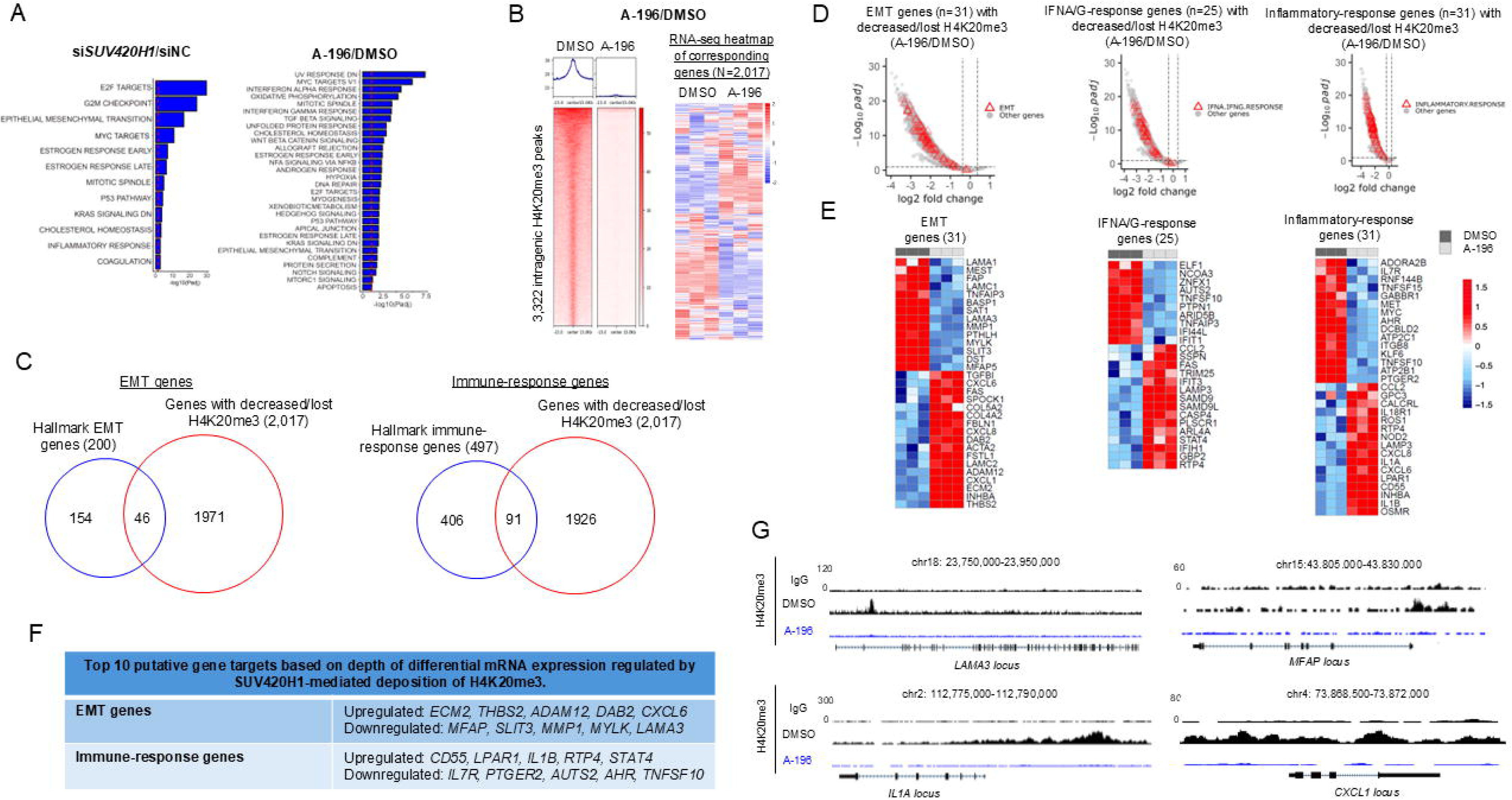
SUV420H1 regulates the expression of specific EMT and immune-response genes through H4K20me3. **(A)** GSEA analysis of significantly enriched Hallmark pathways in HN-SCC-151 cells treated with si*SUV420H1*(#1)/siNC (left) or A-196/DMSO (right) for 6 days. **(B)** Genomic coordinate heatmap of decreased/lost intragenic H4K20me3 peaks (n=3,322) in HN-SCC-151 cells treated with A-196 vs DMSO for 6 days. FDR<0.1, log2FC>0.38. Concordant RNA-seq heatmap of 2,017 corresponding genes with decreased/lost intragenic H4K20me3 peaks and available mRNA expression data; FDR<0.1. **(C)** Venn diagram of overlapping EMT and immune-response (IFNA/G-, inflammatory-response) Hallmark pathway genes and the 2,017 genes with decreased/lost H4K20me3 peaks and detectable mRNA expression (A-196 CUT&RUN and RNA-seq datasets). **(D)** Volcano plots of H4K20me3 peaks overlapping with EMT (127 peaks) or IFNA/G-(100 peaks) and inflammatory-response genes (122 peaks) (promoter and gene body regions only). FDR<0.1, LFC>0.38. **(E)** RNA-seq heatmaps of EMT, IFNA/G-response and inflammatory-response genes with decreased/lost H4K20me3 peaks and differential mRNA expression changes (FDR<0.1). **(F)** Top 10 putative EMT and immune-response gene targets with the “deepest” mRNA expression changes regulated by SUV420H1-mediated H4K20me3. **(G)** UCSC tracks for H4K20me3 obtained with CUT&RUN for H4K20me3 in HN-SCC-151 cells treated with DMSO or A-196 for 6 days. Representative EMT and immune-response genes with decreased/lost H4K20me3 after treatment with A-196 are shown.

Subsequently, the parental and the two *Suv420h1* KO MOC1 cell lines were implanted in the right flank of C57BL/6 mice and tumor growth was monitored. *Suv420h1* KO MOC1 tumors grew significantly less compared to the parental MOC1 tumors (**Fig. 4B**, top and bottom, **Supplementary Fig. 5**). To interrogate the effect of Suv420h1 depletion on the TIME, flank parental and *Suv420h1* KO MOC1 tumors were implanted, surgically resected (day 28 post-tumor implantation), digested into single cells and multicolor flow cytometry was conducted in the parental (NT, n=4-5) and *Suv420h1* KO MOC1 tumors (KO-1, n=4; KO-2, n=3) (**Fig. 4C, Supplementary Fig. 5**). CD8+ T-cells and T-cells showed a trend for an increase in *Suv420h1* KO MOC1 tumors compared to control tumors, however this did not reach significance (**Fig. 4C**). Similarly, the percentage of PD-1+ CD8+ T-cells and PD-L1+ MOC1 tumor cells, as well as the percentage of MOC1 tumor cells that were positive for major histocompatibility complex (MHC) class I H-2Kb/Db was increased but not at a significant level (**Fig. 4C**). Interestingly, the macrophage population was more predominant and increased from ∼10% in the control to ∼30% in the *Suv420h1* KO tumors, while the granulocytic myeloid-derived suppressor cells (gMDSC) were significantly decreased from ∼50% in the control to ∼10-30% in the *Suv420h1* KO tumors (**Fig. 4C**). The monocytic MDSCs (mMDSCs) were not affected (**Fig. 4C**). Furthermore, CD4+ T-cells and B-cells were significantly increased in *Suv420h1* KO MOC1 tumors, though this increase was numerically small (CD4+ T-cells: ∼2% in control to ∼5-7% in *Suv420h1* KO tumors; B-cells: ∼1% in control to 2% in *Suv420h1* KO tumors) (**Fig. 4C**). These data show that Suv420h1 depletion in MOC1 tumors induces a significant increase in the percentage of intratumoral macrophages with a concurrent decrease in the percentage of gMDSCs, while not significantly affecting the percentage of intratumoral CD8+ T-cells. This supports that Suv420h1 may promote innate anti-tumor immunity by altering the balance and/or abundance of macrophages and gMDSCs.

We then evaluated whether Suv420h1 depletion could synergize with anti-PD-1 therapy in this syngeneic mouse model of flank MOC1 tumors. *Suv420h1* KO and NT control MOC1 flank tumors were implanted as described above and mice were treated systemically with anti-PD-1 therapy in the following groups: NT (N=8), NT + anti-PD-1 (N=8), KO-1 (N=7), KO-1 + anti-PD-1 (N=9). Tumor growth and overall survival were monitored for up to 80 days. While anti-PD-1 therapy alone did not restrict tumor growth or prolong survival (**Fig. 4D, E**), the combination of *Suv420h1* KO and anti-PD-1 therapy significantly restricted tumor growth (p<0.001) and prolonged survival (Log-rank, p=0.005) compared to anti-PD-1 therapy or *Suv420h1* KO alone (**Fig. 4D, E**). These findings support that Suv420h1 depletion sensitizes MOC1 tumors to anti-PD-1 therapy.

### SUV420H1 affects proliferation, EMT and immune response pathways in HPV-negative HNSCC cells

To identify pathways regulated by SUV420H1 in HPV-negative HNSCC cells, Gene Set Enrichment Analysis (GSEA) was conducted utilizing the RNA-seq datasets generated in HN-SCC-151 cells after siRNA-mediated depletion (si1, 2, 3 for *SUV420H1*) or enzymatic inhibition (A-196) of SUV420H1 for 6 days. In the si*SUV420H1* RNA-seq datasets, GSEA revealed enrichment of cell cycle and proliferation (E2F targets, G2M checkpoint, MYC targets, Mitotic spindle), EMT and immune response (IFNA-, IFNG-, inflammatory-response) Hallmark pathways (**Fig. 5A, Supplementary Fig. 6, Supplementary Table 2A**). Similar results were obtained with GSEA analysis of the A-196-treated HN-SCC-151 cells, however, the level of enrichment for statistical significance was lower (-log10(Padj)=1-7.5) compared to the level of enrichment observed with the si*SUV420H1* RNA-seq datasets (-log10(Padj)=1-30) (**Fig. 5A, Supplementary Table 2B**). Furthermore, the IFN-response pathways were amongst the top 5 most significantly enriched pathways in the A-196 GSEA analysis, suggesting that the effects of Suv420h1 KO observed in the TIME of MOC1 syngeneic mouse models may be enzymatically mediated. These results are in accordance with the GSEA analysis in the HPV-negative HNSCC and LUSCC cohort of the TCGA (**Fig. 1C**), our observed phenotypes of decreased proliferative potential and invasive capacity with human SUV420H1 depletion in vitro (**Fig. 2B, C, Fig. 3A, B, D**), enhanced immune response with mouse Suv420h1 depletion in vivo (**Fig. 4**), as well as with decreased invasive capacity with human SUV420H1 inhibition (**Fig. 3. D, E**).

### SUV420H1 affects global levels and the genomic distribution of H4K20me3 in HPV-negative HNSCC cells

SUV420H1 is a chromatin modifier that writes H4K20me3 (4, 7). As previously shown above, treatment of three HPV-negative HNSCC cell lines with the enzymatic SUV420H1 inhibitor A-196 lead to an approximately 50-80% reduction in the global levels of H4K20me3 (**Fig. 2D**), supporting that SUV420H1 is a major regulator of H4K20me3 in HPV-negative HNSCC cells. These results were further validated with siRNA-mediated depletion of *SUV420H1* (**Supplementary Fig.7**).

We then sought to assess the genome-wide distribution of SUV420H1 and its effect on the genomic distribution of H4K20me3 using CUT&RUN assays, and to identify putative gene targets that are transcriptionally regulated directly by SUV420H1 and/or by SUV420H1-mediated deposition of H4K20me3. We expected that gene targets bound by SUV420H1 would include genes regulated directly by SUV420H1 in a catalytically-independent manner, as well as genes regulated in a catalytically-dependent manner through SUV420H1-mediated deposition of H4K20me3. While we attempted to conduct CUT&RUN assays for endogenously expressed SUV420H1 and for HA-tagged SUV420H1, no interpretable CUT&RUN signal was obtained through these attempts. We thus focused on identifying putative gene targets of SUV420H1 that are transcriptionally regulated by SUV420H1-mediated H4K20me3 deposition and would thus mediate catalytically-dependent phenotypes, specifically the EMT and possibly the immunomodulation phenotype.

To this purpose, HN-SCC-151 cells were treated with the SUV420H1 inhibitor A-196 (2.5uM) for 6 days, and CUT&RUN was conducted for H4K20me3. The same experiment was performed with HN-SCC-151 cells treated with *SUV420H1*-targeting siRNAs (si1) versus control siNC to validate the results obtained with the A-196 inhibitor. 10,196 H4K20me3 and 16,503 H4K20me3 peaks were called in HN-SCC-151 cells in the DMSO or siNC treated state. SUV420H1 enzymatic inhibition or siRNA-mediated depletion decreased the genome-wide deposition of H4K20me3, indicated by the significantly decreased average normalized tag counts corresponding to H4K20me3-binding sites throughout the genome (**Supplementary Fig. 8A**). Nearly 99% of all H4K20me3 peaks were decreased or lost after A-196 or si*SUV420H1* treatment, supporting that SUV420H1 is the primary regulator of H4K20me3 genomic deposition in HPV-negative HNSCC cells (**Supplementary Fig. 8B**, C). In both systems, approximately 65% of the decreased/lost H4K20me3 peaks were intragenic, while roughly 35% had an intergenic distribution, suggesting a role for SUV420H1-mediated H4K20me3 deposition predominantly in the direct regulation of genes, and less in the regulation of intergenic regulatory elements (**Supplementary Fig. 8C**).

These results show that SUV420H1 is the primary regulator of H4K20me3 genomic deposition in HPV-negative HNSCC cells.

### SUV420H1 regulates the expression of EMT and immune-response genes through H4K20me3

To identify genes directly regulated by SUV420H1-mediated H4K20me3 deposition, we focused our analysis on genes with decreased/lost intragenic H4K20me3 in the A-196 CUT&RUN dataset (HN-SCC-151 cells). 3,322 intragenic peaks were called in DMSO-treated HN-SCC-151 cells, and these corresponded to 2,017 genes with available mRNA expression data (**Fig. 5B**) (**Supplementary Table 3**). A significant overlap of 83% of genes with decreased/lost H4K20me3 intragenic peaks was observed between the A-196 and the si*SUV420H1* CUT&RUN datasets, providing higher confidence for the identified candidate genes as genes regulated by SUV420H1-mediated H4K20me3 (**Supplementary Fig. 8D**). Interestingly, of the 2,017 genes identified in the A-196 datasets, the decrease/loss of H4K20me3 was associated with transcriptional upregulation in ∼50% of genes, consistent with the canonical repressive function of H4K20me3 (**Fig. 5B**). However, ∼35% of genes showed transcriptional downregulation, supporting a non-canonical function of SUV420H1 and H4K20me3 as activators of these genes (FDR<0.1) (**Fig. 5B**). A similar transcriptional pattern for genes with decreased/lost H4K20me3 was obtained in the RNA-seq dataset of HN-SCC-151 cells treated with si*SUV420H1* depletion, with 52% of genes upregulated and 38% of genes downregulated (**Supplementary Fig. 8E**).

To identify specific genes directly regulated by SUV420H1-mediated H4K20me3 deposition that could putatively dictate the observed catalytically-dependent phenotypes of EMT and possibly immunomodulation, the gene sets of the EMT and immune-response (IFNA-, IFNG- and inflammatory-response) Hallmark pathways that were found enriched in the A-196 RNA-seq GSEA analysis (**Fig. 5A, Supplementary Table 4**) were overlapped with the genes with decreased/lost H4K20me3 peaks in the A-196 CUT&RUN dataset with available mRNA expression data (**Supplementary Table 3**). 46 EMT and 91 IFNA/G/inflammatory-response genes with decreased/lost intragenic H4K20me3 peaks were identified (**Fig. 5C, Supplementary Table 5**). We then further focused on the genes with differential mRNA expression induced by the treatment with A-196 (FDR<0.1). With this approach, 31 EMT and 56 IFNA-/IFNG-/inflammatory-response genes were identified as putative gene targets directly regulated by SUV420H1-mediated deposition of H4K20me3 (**Fig. 5D, E, Supplementary Table 6A, B**). **Fig. 5F** summarizes the top 10 EMT and immune-response genes with the “deepest” differential mRNA expression changes.

Of these EMT genes, microfibrillar-associated protein (*MFAP*), matrix metallopeptidase 1 (*MMP1*) and laminin subunit alpha 3 (*LAMA3*) which promote invasion, were significantly downregulated with A-196 treatment (**Fig. 5F, Supplementary Table 6A**). Examples of genome-wide mapping tracks of some of these EMT genes are shown in **Fig. 5G**. We further interrogated all differentially expressed EMT genes regardless of intragenic decrease/loss of H4K20me3 (**Supplementary Table 7A**) and found that collagenase type V alpha 3 chain (*COL5A3*), fibrillin 1 (*FBN1*), fibronectin 1 (*FN1*), integrin subunit beta 1 (*ITGB1*) and laminin subunit alpha 1 (*LAMA1*), amongst other EMT promoting genes, were significantly downregulated with A-196 treatment.

Of the immune-response genes with decreased/lost intragenic H4K20me3, the receptor transporter protein 4 (*RTP4*), signal transducer and activator of transcription 4 (*STAT4*), sterile alpha motif domain-containing protein 9 ligand (*SAMD9L*), lysosome-associated membrane protein 3 (*LAMP3*) and interferon-induced protein with tetratricopeptide repeats 3 (*IFIT3*), amongst other IFNA/G-response genes, were found significantly upregulated with A-196 treatment (**Supplementary Table 6B**). Interrogating all differentially expressed immune-response genes regardless of intragenic decrease/loss of H4K20me3 (**Supplementary Table 7B**), human leukocyte antigens-A, -B and -C (*HLA-A*, *HLA-B*, *HLA-C*), and proteasome subunit beta types 8, 9 and 10 (*PSMB8*, *PSMB9*, *PSMB10*), which code for major components of the antigen presentation process, were also found significantly upregulated.

These EMT and immune-response genes may be putative culprits for the invasive and immunomodulatory phenotypes associated with SUV420H1-mediated H4K20me3.

### SUV420H1-mediated deposition of H4K20me3 regulates the expression of myeloid-attracting chemokines

Interestingly, within the list of immune-response genes with decreased/lost intragenic H4K20me3 and differential mRNA expression changes (**Fig. 5F, Supplementary Fig. 6B**), we identified a number of genes known to promote trafficking, migration and activation of macrophages to sites of inflammation, as we observed in the TIME of *Suv420h1* KO MOC1 tumors. Specifically, interleukin-1 beta (*IL1B*) and lysophosphatidic acid receptor 1 (*LPAR1*) were significantly upregulated by A-196 treatment. Furthermore, interleukin-1 alpha (*IL1A*), and C-X-C motif chemokine ligand 1 (*CXCL1*) and C-X-C motif chemokine ligand 8 (*CXCL8*), which also promote the trafficking of macrophages to sites of inflammation, were also significantly upregulated (**Supplementary Table 6 A, B**). Examples of genome-wide mapping tracks of some of these genes are shown in **Fig. 5G**. We then further interrogated all differentially expressed immune-response genes regardless of intragenic decrease/loss of H4K20me3 (**Supplementary Table 7B**) and found that interleukin-15 (*IL15*), which promotes macrophage trafficking, was also significantly upregulated. Importantly, interleukin-6 (*IL6*), which codes for a neutrophil- and MDSC-attracting cytokine, was identified as the most downregulated gene amongst all immune-response genes regardless of intragenic decrease/loss of H4K20me3 (**Supplementary Table 7B**). C-X-C motif chemokine ligand 17 (*CXCL17*), a chemokine that has been reported to induce gMDSC intratumoral trafficking was also significantly downregulated after A-196 treatment.

These results support that genes coding for myeloid-attracting chemokines, such as *IL1A*, *IL1B*, *LPAR1*, *CXCL1*, *CXCL8*, *IL15*, *IL6* and *CXCL17*, are regulated by SUV420H1-mediated deposition of H4K20me3, and could be the culprits inducing the increased macrophage and decreased gMDSC infiltration observed in the TIME of *Suv420h1* KO MOC1 tumors.

## DISCUSSION

Our findings reveal catalytically-dependent and -independent roles for SUV420H1 in HPV-negative HNSCC: a catalytically-dependent role in regulating invasion and possibly an immunosuppressive TIME through H4K20me3, and a catalytically-independent role in regulating cancer cell proliferation and cell cycle progression. Through a combination of genome-wide profiling using CUT&RUN assays, RNA-seq and in vivo approaches, we demonstrate that SUV420H1 acts as a chromatin modifier with oncogenic and immunomodulatory activity in HPV-negative HNSCC.

Using the TCGA genomic dataset, we recently identified SUV420H1 as one of the most frequently recurrently amplified protein methyltransferases in HPV-negative HNSCC (14), with ∼36% of tumors exhibiting gain or focal amplification of *SUV420H1* (**Supplementary Fig.1**). *SUV420H1* gain and amplification was associated with increased mRNA expression (**Fig.1 A, B**), which correlated with worse overall survival in HPV-negative HNSCC patients (**Fig. 1D**). The high prevalence of gain/amplification of *SUV420H1* was also observed in other squamous carcinomas, including lung and bladder SCC, supporting that SUV420H1 may function as a pan-squamous epigenetic driver (**Supplementary Fig .1**).

Mechanistically, we showed that SUV420H1 depletion halts the proliferative and colony-forming capacity of HPV-negative HNSCC cells through a catalytically-independent manner. Specifically, siRNA-mediated *SUV420H1* depletion resulted in near-complete suppression of proliferation and induced G2/M cell cycle arrest (**Fig. 2B, C, Fig. 3A, B**). In contrast, neither pharmacologic inhibition of SUV420H1’s catalytic activity using the A-196 SUV420H1-inhibitor, nor expression of mutant, catalytically-inactive SUV420H1 recapitulated this phenotype, despite reduced global H4K20me3 levels (**Fig. 2D-J, Fig. 3A-C**). These results support a catalytically-independent function of SUV420H1 in regulating proliferation, potentially through chromatin scaffolding or recruitment of chromatin co-factors (5, 8). In contrast to its catalytically-independent role in proliferation, our findings demonstrated that SUV420H1 regulates the invasive capacity of HPV-negative HNSCC cells through a catalytically-dependent mechanism. *SUV420H1* depletion through siRNAs, enzymatic inhibition, and expression of catalytically-inactive SUV420H1 significantly impaired invasion (**Fig 3D, E**). This paradigm aligns with emerging evidence of non-canonical, catalytically-independent functions of other histone methyltransferases, including Enhancer of Zeste 2 Polycomb Repressive Complex 2 Subunit (EZH2) and SET domain containing 2 (SETD2), which regulate DNA repair, transcription, and replication licensing through protein-protein interactions rather than histone methylation (17–19).

Our transcriptomic analysis revealed consistent enrichment of IFNA/G-response pathways in both TCGA tumors and our in vitro RNA-seq datasets of A-196- and si*SUV420H1*-treated HPV-negative HNSCC cells (**Fig. 1C, Fig. 5A**). Furthermore, we observed a decrease/loss of intragenic H4K20me3 and increased mRNA expression of a number of IFNA/G-response genes in the A-196 CUT&RUN and RNA-seq datasets (**Fig. 5B-G**). These results are in accordance with the reported role of H4K20me3 in the silencing of IFN-response genes. Specifically, recently published work from our group demonstrated that SET-and MYND-domain-containing 3 (SMYD3), a protein methyltransferase, maintains repression of IFN-response genes through intragenic H4K20me3 deposition, and that SMYD3 depletion leads to re-expression of IFN-response genes, including *CXCL9*/*10* and *11* CD8+ T-cell attracting chemokine genes, increased intratumoral CD8+ T-cell infiltration and sensitization of MOC1 tumors to anti-PD-1 therapy (20). Given that SUV420H1 is the principal methyltransferase for H4K20me3, it is plausible that SUV420H1 functions in concert with, or in parallel to SMYD3 to silence IFN-response genes through H4K20me3, a hypothesis that merits further investigation. Ramponi et al. also recently demonstrated that SUV420H1/2-mediated H4K20me3 directly represses IFN-response gene expression at or near promoters in persister cancer cells, and that SUV420H1 depletion leads to robust activation of IFN-response programs in human melanoma, non-small cell lung cancer and breast cancer cell lines (21). Of significance, H4K20me3 has been reported to maintain transcriptional repression of endogenous retroviral elements and retrotransposons, such as LINE-1, and its loss induces derepression of these repetitive elements, induction of cytosolic sensing of double-stranded nucleic acids by cGAS–STING and RIG-I–MAVS pathways, with subsequent activation of IFN-response pathways (8, 22–26). While *STING1* was significantly upregulated in the si*SUV420H1* but not the A-196 RNA-seq dataset (**Supplementary Table 2**), the possibility that SUV420H1 may also silence retransposon sequences through H4K20me3 further repressing IFN responses in HPV-negative HNSCC cells would need to be further and more systematically interrogated through whole RNA-sequencing experiments.

Additionally, using a syngeneic MOC1 mouse model of HPV-negative HNSCC, we demonstrated that Suv420h1 modulates the TIME in vivo (**Fig. 4**). CRISPR-mediated *Suv420h1* knockout in MOC1 mouse tumors significantly attenuated tumor growth (**Fig. 4B**) and triggered immune cell reconstitution of the TIME. Notably, *Suv420h1* loss significantly increased the intratumoral macrophage compartment with an associated reduction of gMDSCs, and a modest but non-significant increase in the CD8+ T cell population (**Fig. 4C**). Concordantly, SUV420H1 inhibition with A-196 in human HN-SCC-151 cells in vitro induced loss of intragenic H4K20me3 deposition and concurrent transcriptional upregulation of *IL1A*, *IL1B*, *LPAR1*, *CXCL1*, *CXCL8*, as well as secondary upregulation of *IL15*, which code for chemokines that induce macrophage attraction (27–28). *IL6* and *CXCL17* did not demonstrate a decrease/loss of intragenic H4K20me3, but they were significantly downregulated probably as indirect gene targets of SUV420H1, which could explain the decrease in the intratumoral gMDSC infiltration in *Suv420h1* KO MOC1 tumors (27, 29). *CXCL10*, which codes for a CD8+ T-cell attracting chemokine, was upregulated in the si*SUV420H1* but not the A-196 RNA-seq dataset. Interestingly, we previously reported that intratumoral depletion of Smyd3 using anti-sense oligonucleotides (ASOs) also increased the CD8+ T-cells and macrophages, but did not affect the gMDSC population of MOC1 tumors (20). In contrast, Suv420h1 depletion seemed to impart a more specific effect on the myeloid compartment of MOC1 tumors. While the contrast of these results begets great caution given that they are drawn from different experiments, it is possible that SUV420H1 may have a more specific effect on macrophage and gMDSC trafficking. The aforementioned results suggest that SUV420H1 promotes an immunosuppressive TIME, possibly through transcriptional regulation of myeloid-attracting chemokine/cytokine genes, such as *IL1A*, *IL1B*, *LPAR1*, *CXCL1*, *CXCL8*, *IL15*, *IL6* and *CXCL17*, which alter the balance of anti-tumor macrophages/pro-tumorigenic MDSCs towards a pro-tumorigenic TIME.

Importantly, despite the only modest and non-significant increase in intratumoral CD8+ T-cells and a predominant effect on the myeloid compartment, *Suv420h1* KO MOC1 tumors exhibited enhanced sensitivity to anti-PD-1 therapy, further potentiating the anti-tumor efficacy of *Suv42h0h1* KO (**Fig. 4D**). Several reports have documented the immunosuppressive role of tumor-associated macrophages (TAMs) in HPV-negative HNSCC and other squamous cancers, as well as their contribution to resistance against PD-1/PD-L1 blockade (30, 31). Interestingly, PD-1 expression on macrophages has been associated with impaired phagocytosis (32), and PD-1/PD-L1 inhibition in preclinical models has been shown to restore macrophage-mediated antigen presentation and promote tumor clearance (33). This suggests that SUV420H1 depletion in HPV-negative HNSCC tumors may synergize with anti-PD-1 immune checkpoint blockade by (i) inducing the re-expression of IFN-response genes, (ii) increasing the intratumoral influx of macrophages and decreasing that of gMDSCs towards a more immunostimulatory TIME through transcriptional regulation of myeloid-attracting chemokines/cytokine genes *IL1A*, *IL1B*, *LPAR1*, *CXCL1*, *CXCL8*, *IL15* and *IL6*, and (iii) by promoting the anti-tumorigenic function of macrophages.

Regarding the effect of SUV420H1 on the genome-wide deposition of H4K20me3 in HPV-negative HNSCC cells, CUT&RUN profiling in A-196-treated HPV-negative HNSCC cells revealed a genome-wide loss of predominantly intragenic H4K20me3 peaks, implicating SUV420H1 as the primary regulator of H4K20me3 deposition in HPV-negative HNSCC. Interestingly, integration of the H4K20me3 genome-wide mapping with RNA-seq showed that, while ∼40% of genes with intragenic reduction/loss of H4K20me3 were transcriptionally upregulated, consistent with the established repressive function of H4K20me3, another ∼40% of genes with intragenic reduction/loss of H4K20me3 were downregulated, suggesting a non-canonical, activating role for H4K20me3 for specific gene sets. This interesting finding implies a “bifaceted” function for H4K20me3, either as a repressive or an activating histone mark. Such non-canonical “bifaceted” behaviors have been reported with certain chromatin modifiers (20, 34, 35), however the specific mechanisms that could render this “bifaceted” behavior for H4K20me3 remain elusive and merit further investigation.

Our data support that SUV420H1 promotes tumor growth through catalytically-independent mechanisms, but modulates invasion and also possibly immunosuppression via its catalytic activity. These findings have major implications for therapeutic targeting. While the SUV420H1 inhibitor A-196 effectively depleted its enzymatic endproduct H4K20me3 and impaired invasion, it failed to block SUV420H1-dependent proliferation and cell cycling. Thus, catalytic inhibitors alone may be therapeutically insufficient, and strategies that degrade or disrupt the non-catalytic functions of SUV420H1 may be required. Proteolysis-targeting chimeras (PROTACs), anti-sense oligonucleotides (ASOs) or small molecules that interfere with protein-protein interactions of SUV420H1 likely represent more effective therapeutic approaches (34, 36–38).

Our study has a number of shortcomings. Although we were able to map genome-wide changes in H4K20me3, our attempts to perform CUT&RUN using antibodies against endogenous or HA-tagged SUV420H1 yielded poor signals, limiting our ability to identify direct gene targets of SUV420H1 that could explain its catalytically-independent role. Additionally, while A-196 is a potent SUV420H1/2 inhibitor, its dual specificity complicates attribution of effects solely to SUV420H1. However, the consistency of results across *SUV420H1*-mutant and A-196-treated HNSCC cell lines suggests that our results with the A-196 inhibitor are specific to SUV420H1. Regarding the effect of Suv420h1 on the TIME, while we identified a possible role of macrophages in the anti-tumor efficacy and synergy of *Suv420h1* KO with anti-PD-1, macrophage depletion experiments in *Suv420h1* KO tumors treated with anti-PD-1 will be necessary to validate the functional role of macrophages in this context. Along the same lines, the effect of SUV420H1 inhibition on the expression of myeloid-attracting chemokine/cytokine genes requires further validation at the protein level. Furthermore, it would be necessary to also assess the direct effects of Suv420h1 inhibition/depletion on macrophage reprogramming through functional analyses of macrophage activation and antigen presentation and through in vivo experiments with syngeneic mice systemically treated with Suv420h1 inhibitors or degraders. Also, given the reported role of SUV420H1 in DNA repair (7) and with chemoradiotherapy constituting a standard of care therapeutic approach in HPV-negative HNSCC, interrogating the effect of SUV420H1 on chemoradiotherapy resistance merits further investigation. Finally, our finding that SUV420H1-mediated H4K20me3 appears to function not only as a repressive mark on certain gene sets, but also as an activating mark on other gene sets within the same cell context is a provocative result that requires further mechanistic interrogation.

Taken together, our findings support a model wherein SUV420H1 promotes HPV-negative HNSCC progression through complementary catalytically-independent and -dependent functions. In the nucleus, SUV420H1 facilitates proliferation and cell cycle progression independently of H4K20me3, while catalytically depositing H4K20me3 at EMT and immune-related genes to enhance invasion and shape an immunosuppressive TIME (**Fig. 6**). These results nominate SUV420H1 as an oncogenic chromatin modifier in HPV-negative HNSCC and provide rationale for its depletion/degradation as a novel therapeutic approach to hinder proliferation, invasion and potentiate sensitization to anti-PD-1 immunotherapy in patients with HPV-negative HNSCC. The presence of *SUV420H1* gain/amplification may represent a biomarker to rationally select patients with HPV-negative HNSCC that are vulnerable to SUV420H1 therapeutic targeting.

## ACKNOWLEDGEMENTS

This research was supported [in part] by the Intramural Research Program of the National Institutes of Health (NIH). The contributions of the NIH author(s) were made as part of their official duties as NIH federal employees, are in compliance with agency policy requirements, and are considered Works of the United States Government. However, the findings and conclusions presented in this paper are those of the author(s) and do not necessarily reflect the views of the NIH or the U.S. Department of Health and Human Services. Grammarly AI was utilized to assess for grammatical errors and to enhance conciseness and tone of the manuscript.

## AUTHOR CONTRIBUTIONS

AM and ML were responsible for designing the work that led to the submission, acquiring data, interpreting results, drafting and revising the manuscript. MSD, MP and JA were responsible for acquiring data and interpreting results. SK, MJ and HC were responsible for analyzing large-scale bioinformatics datasets and interpreting results. KM and EE were responsible for acquiring, analyzing and interpreting flow cytometry and IHC based results. VS was responsible for conceiving and designing the work that led to the submission, interpreting results, drafting and revising the manuscript. All authors revised the manuscript, approved the final version and agreed to be accountable for all aspects of the work.

## COMPETING INTERESTS

The authors declare no competing interests.

## DATA AVAILABILITY

Data is provided within the manuscript or supplementary information files. All sequencing raw and processed data will be deposited in the Gene Expression Omnibus (GEO) database and will become publicly available under a specific GEO series GEO upon acceptance of this manuscript for publication.

---

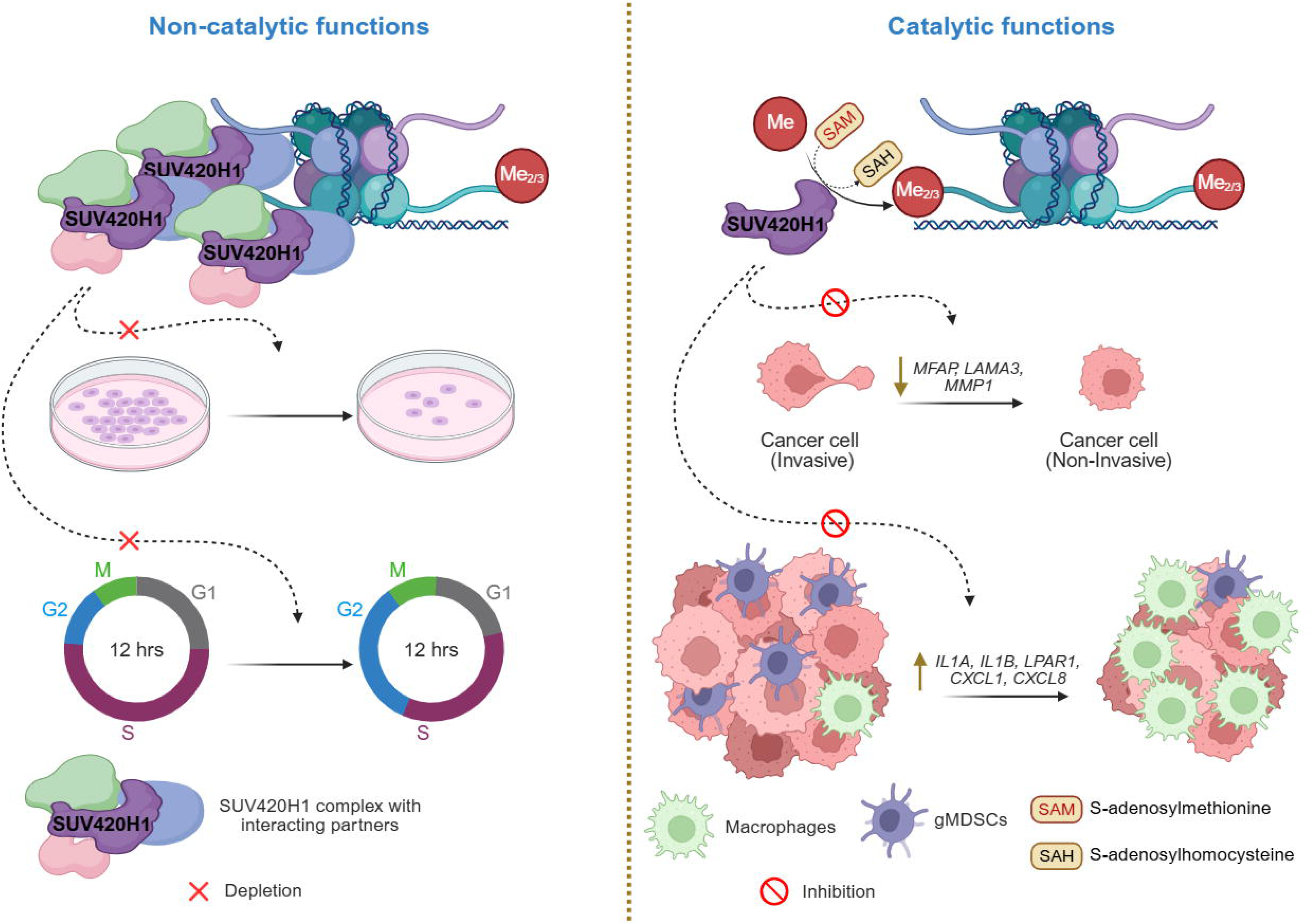

## Notes

### Competing Interest Statement

The authors have declared no competing interest.

