## Supplementary Figures for "Oncogenic and immunomodulatory functions of SUV420H1 in HPV-negative head and neck squamous cell carcinoma"

**Supplementary Data**

**Supplementary Figure 1.** Genetic amplification, gain and mutation rate of *SUV420H1* (*KMT5B*) in HPV-negative head and neck, bladder, lung, cervical and esophageal squamous cell carcinomas (SCC) (TCGA datasets).

| Dataset | Sample Number | Alteration Frequency (%) | | |
|---|---|---|---|---|
| | | Amplification | Gain | Missense Mutation |
| Head and Neck Squamous Cell Carcinoma (TCGA, Firehose Legacy) | 428 | 6.5 | 29.1 | <1 |
| Bladder Urothelial Carcinoma (TCGA, Firehose Legacy) | 413 | 3.9 | 22.8 | 2.2 |
| Lung Squamous Cell Carcinoma (TCGA, Firehose Legacy) | 511 | 2.7 | 25.0 | <1 |
| Cervical Squamous Cell Carcinoma (TCGA, PanCancer Atlas) | 297 | 1.3 | 10.7 | <1 |

**Supplementary Figure 2.** Subcellular localization of SUV420H1 in HPV-negative HNSCC tumors. Immunohistochemistry (IHC) for SUV420H1 was conducted in a tissue microarray (ATA 14-4) of HPV-negative HNSCC tumors (n=22). **(A)** Examples of SUV420H1 IHC in two HPV-negative HNSCC tumors. **(B)** Average protein expression levels (H-score) of SUV420H1 in the nuclear and cytoplasmic compartments of HPV-negative HNSCC tumors assessed by IHC. **(C)** Average protein expression levels (H-score) of SUV420H1 in the tumor compared to stroma compartment of HPV-negative HNSCC sections. Student t-test, *** p<0.001, **** p<0.0001.

**(A)**

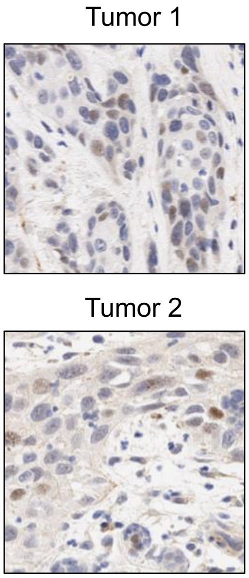


**(B)**

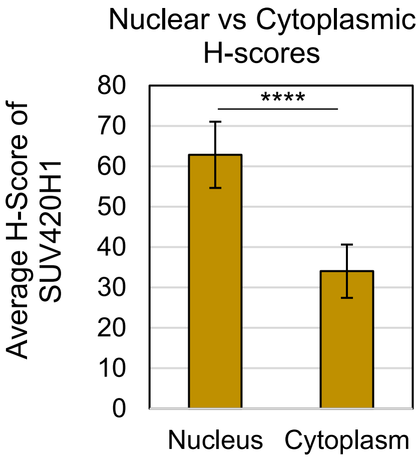


**(C)**

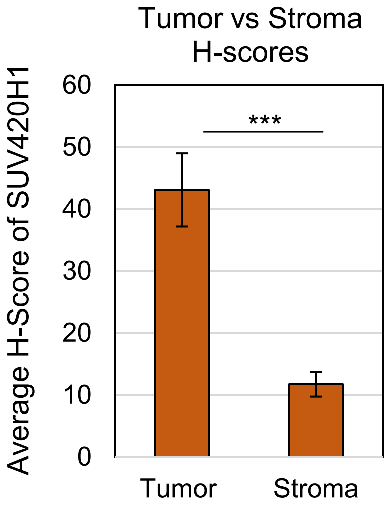

**Supplementary Figure 3.** Gene Set Enrichment Analysis (GSEA) for SUV420H1 in the HPV-negative HNSCC cohort of the Firehose Legacy, TCGA. Significant pathways with padj<0.1 are shown. The GSEA was performed on a ranked gene list. Genes were ranked by their Pearson correlation with *SUV420H1* mRNA. Hallmark pathways shown based on adjusted p-values

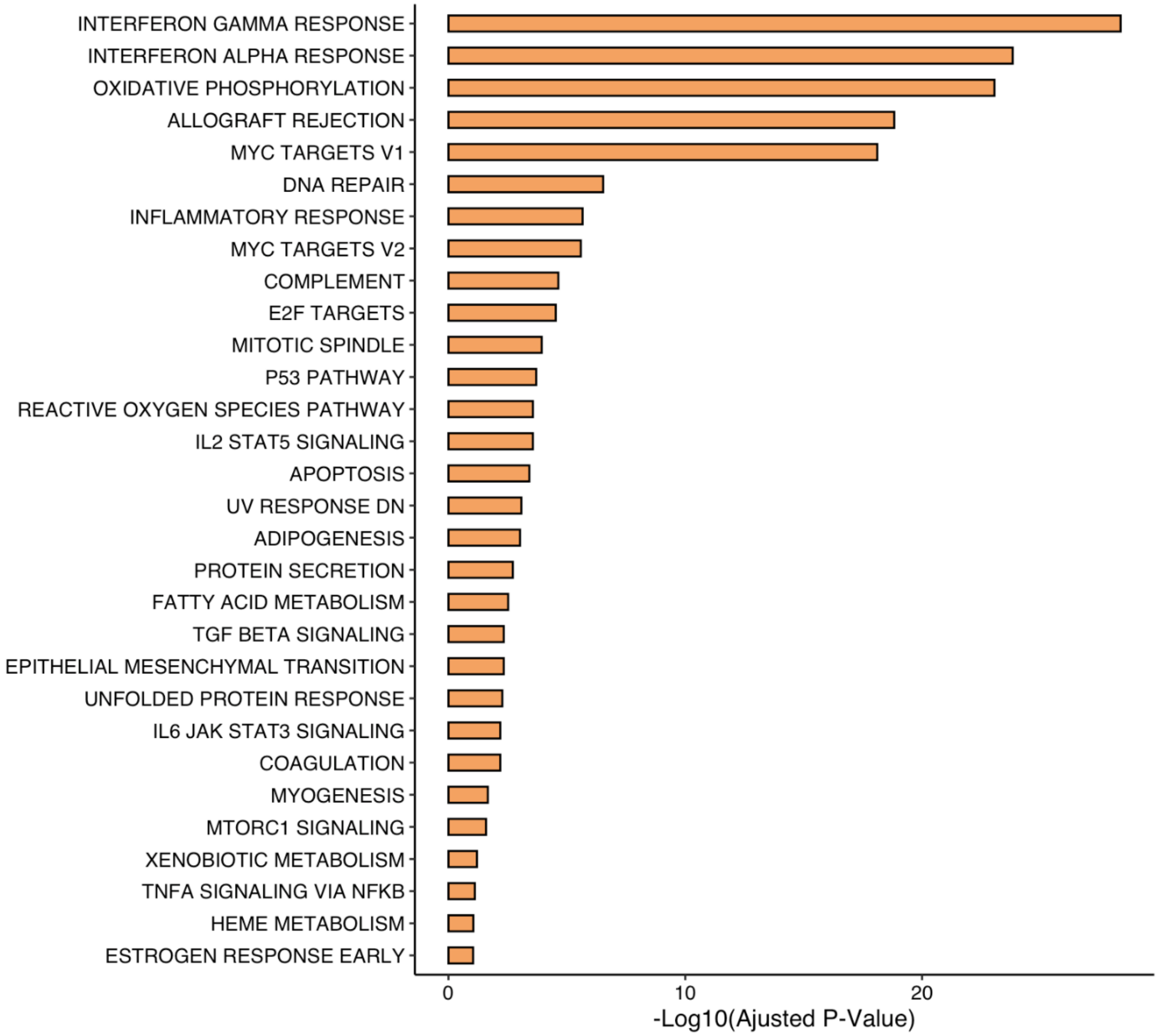

**Supplementary Figure 4.** SUV420H1-targeting siRNAs are specific for *SUV420H1* mRNA but not for *SUV420H2* and *SETD8* H4K20 methyltransferase mRNAs. HN-SCC-151 cells were treated with negative control (siNC) or three different siRNAs targeting *SUV420H1* mRNA (si1, si2, si3) for 72h. Cells were collected, RNA was extracted and qPCR was performed to assess mRNA expression levels of *SUV420H1*, *SUV420H2* and *SETD8*. *GAPDH* mRNA levels were used for normalization. Values represent results from three separate biological replicates per condition. Student t-test, ***p<0.001, ****p<0.0001.

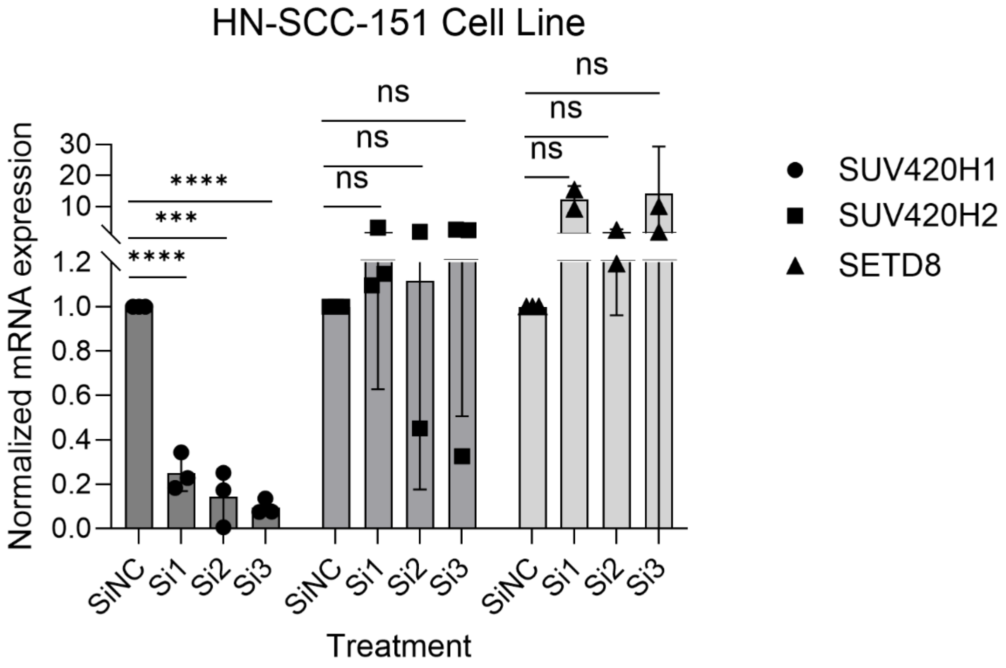

**Supplementary Figure 5.** Mouse experiment of parental (negative control; NT) and *Suv420h1* KO-1 and KO-2 MOC1 tumors used for tumor flow cytometry of Fig. 4C. The right flank of C57BL/6 mice was implanted with 5 million NT (N=5), KO-1 (N=6) or KO-2 (N=6) MOC1 cells. Tumor growth was monitored twice weekly up to day 28, when mice were sacrificed and processed for flow cytometry. Unpaired t-test, day 28, NT versus KO-1, p=0.1, NT versus KO-2, ** p<0.001.

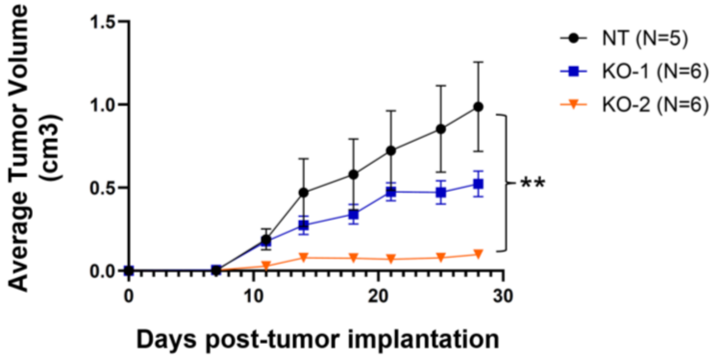


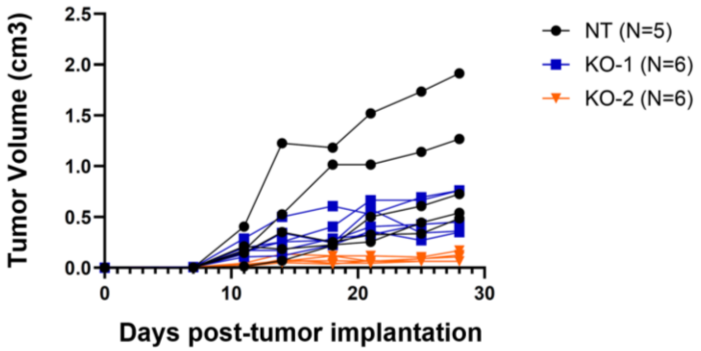

**Supplementary Figure 6.** GSEA analysis of HN-SCC-151 cells treated with si2 and si3 versus control siRNA targeting *SUV420H1*. Padj<0.1, Fold Change (FC) >1.3. H Gene Sets from MSigDB shown.

<u>si*SUV420H1*#2</u>

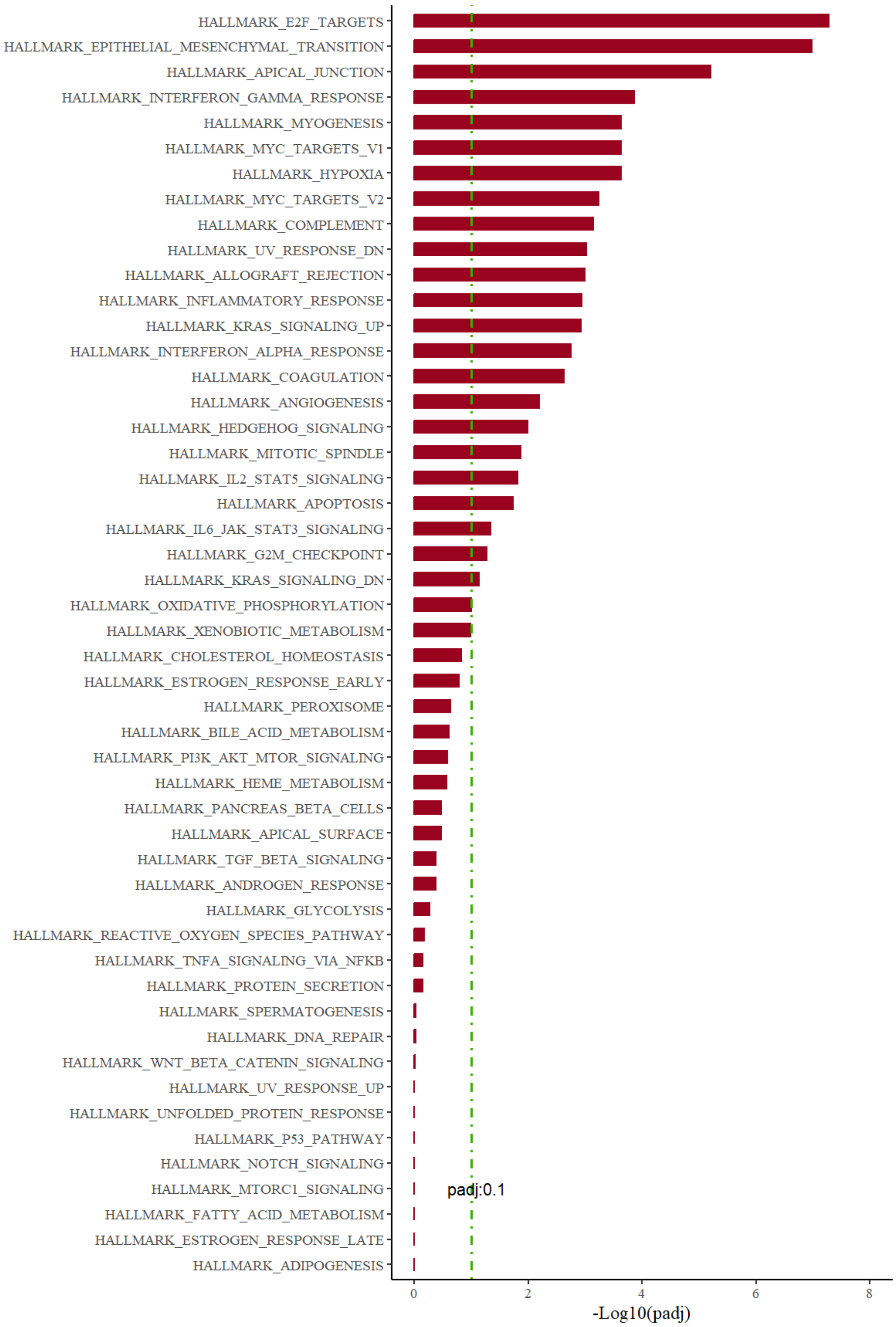

### si*SUV420H1#3*

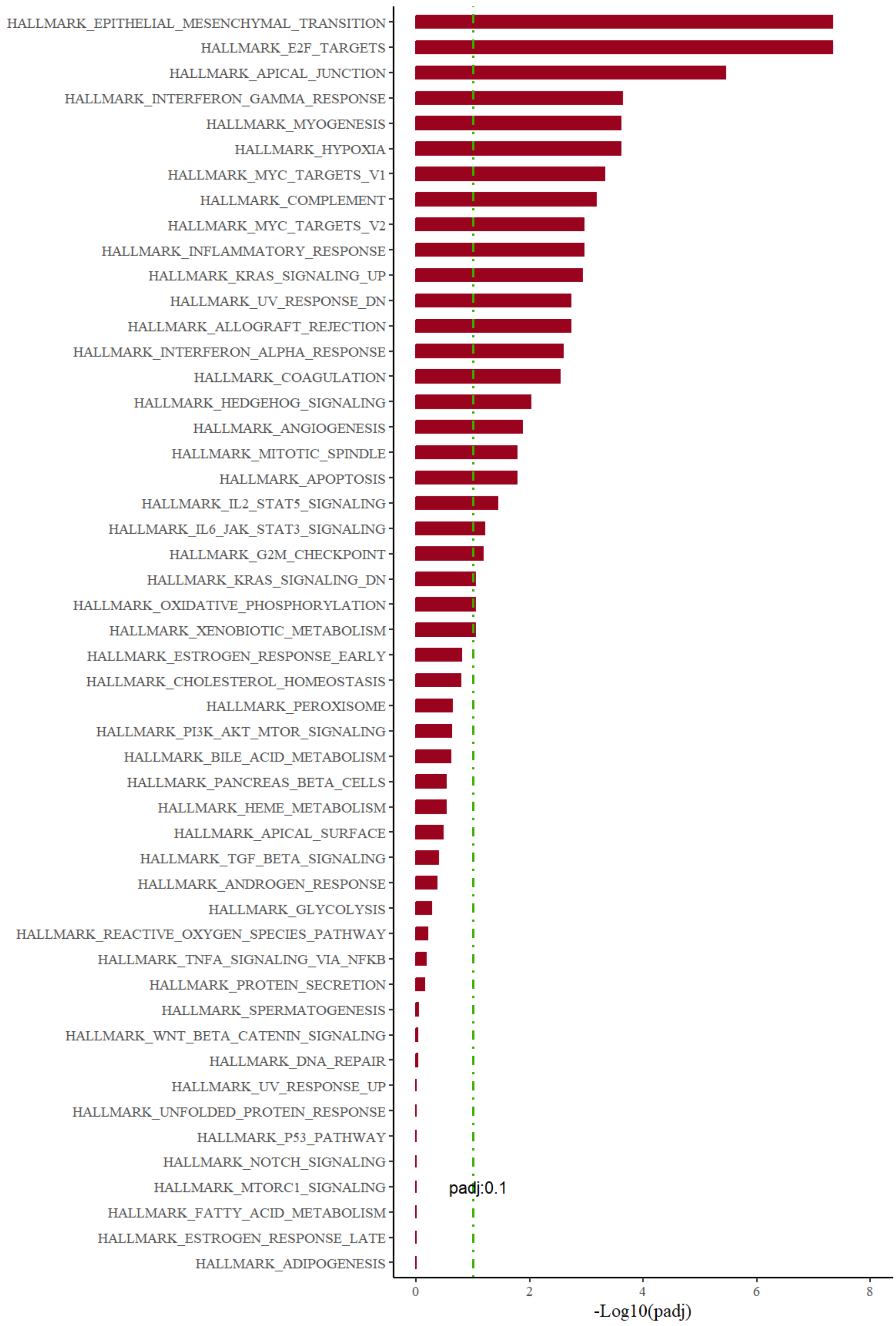

**Supplementary Figure 7.** Global levels of H4K20me3 are decreased after siRNA-mediated *SUV420H1* depletion in HN-6 cells for 6 days. HN-6 cells were treated with *SUV420H1*-targeting (si1, si2, si3) versus control (siNC) siRNAs for 6 days in biological duplicates (for si2 and si3). Cells were then collected and nuclear extraction was conducted. 5ug of nuclear extracts were loaded and blotted for H4K20me3. H3 was used as a loading control. Densitometry graphs are shown at the bottom and represent the mean +/-SEM of the two biological replicates for si1, si2, and si3.

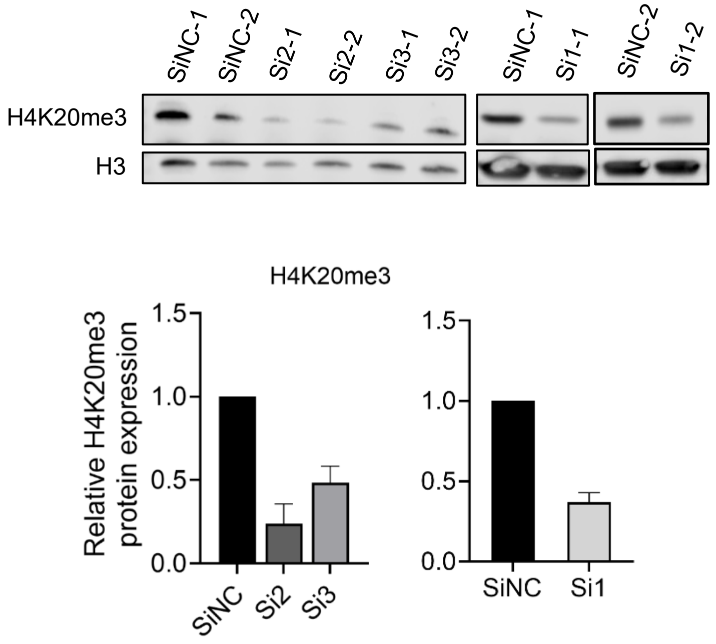

**Supplementary Figure 8.** SUV420H1 is a major regulator of SUV420H1 genomic deposition in HPV-negative HNSCC cells. **(A)** Tag density box plots for H4K20me3 in control (DMSO or siNC) and A-196 (2.5uM) or si*SUV420H1* treated HN-SCC-151 cells for 6 days. **(B)** Volcano plot of all H4K20me3 peaks in HN-SCC-151 cells treated with A-196/DMSO or si*SUV420H1*/siNC for 6 days. FDR<0.1, log2FC>0.38. **(C)** Genome-wide distribution pattern of H4K20me3 in HN-SCC-151 cells before and after A-196 or si*SUV420H1* treatment for 6 days. **(D)** Venn diagram of genes with detectable mRNA expression and intragenic decrease/loss of H4K20me3 in the A-196 and si*SUV420H1* CUT&RUN datasets. **(E)** Genomic coordinate heatmaps of decreased/lost intragenic H4K20me3 peaks (n=10,350) in HN-SCC-151 cells treated with si*SUV420H1* or control siNC for 6 days. FDR<0.1, log2FC>0.38. Concordant RNA-seq heatmap of 2,648 corresponding genes with decreased/lost intragenic H4K20me3 peaks and available RNA-seq data; FDR<0.1, LFC<0.38.

**(A)**

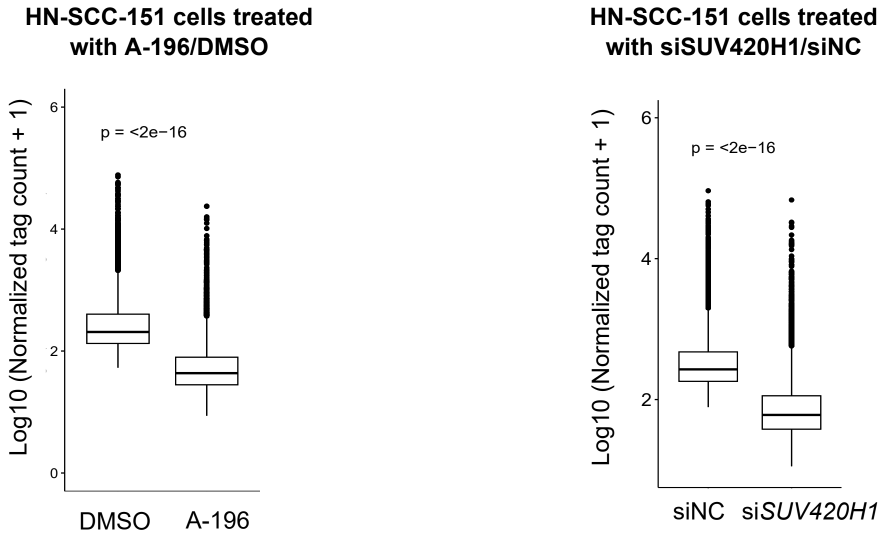


**(B)**

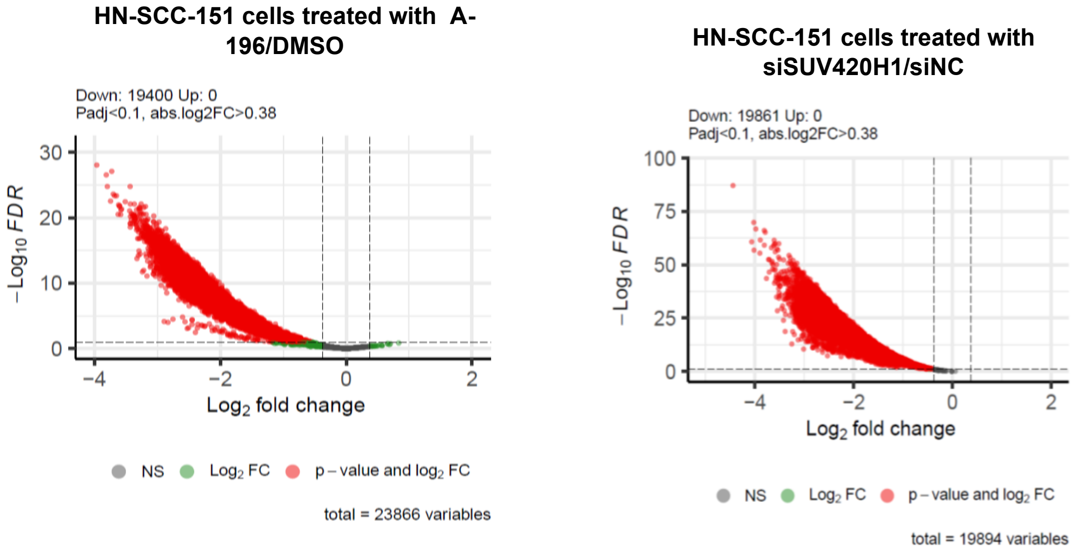

(C)

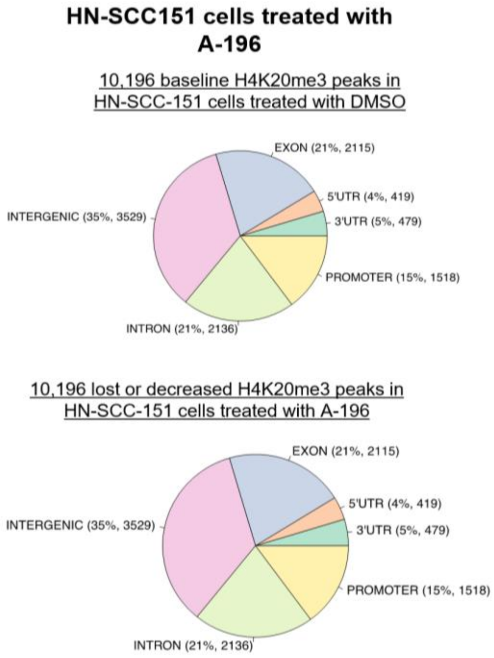

HN-SCC151 cells treated with A-196
10,196 baseline H4K20me3 peaks in HN-SCC-151 cells treated with DMSO
EXON (21%, 2115)
5'UTR (4%, 419)
3'UTR (5%, 479)
PROMOTER (15%, 1518)
INTRON (21%, 2136)
INTERGENIC (35%, 3529)
10,196 lost or decreased H4K20me3 peaks in HN-SCC-151 cells treated with A-196
EXON (21%, 2115)
5'UTR (4%, 419)
3'UTR (5%, 479)
PROMOTER (15%, 1518)
INTRON (21%, 2136)
INTERGENIC (35%, 3529)
HN-SCC151 cells treated with siSUV420H1
16,503 baseline H4K20me3 peaks in HN-SCC-151 cells treated with siNC
EXON (20%, 3356)
5'UTR (3%, 554)
3'UTR (5%, 762)
PROMOTER (10%, 1617)
INTRON (25%, 4068)
INTERGENIC (37%, 6146)
16,494 lost or decreased H4K20me3 peaks in HN-SCC-151 cells treated with siSUV420H1
EXON (20%, 3354)
5'UTR (3%, 553)
3'UTR (5%, 759)
PROMOTER (10%, 1616)
INTRON (25%, 4068)
INTERGENIC (37%, 6144)


(D)

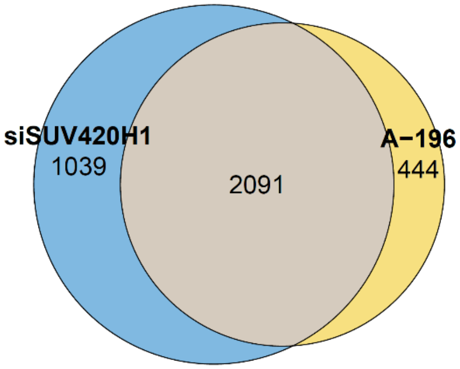

siSUV420H1
1039
2091
A−196
444


(E)

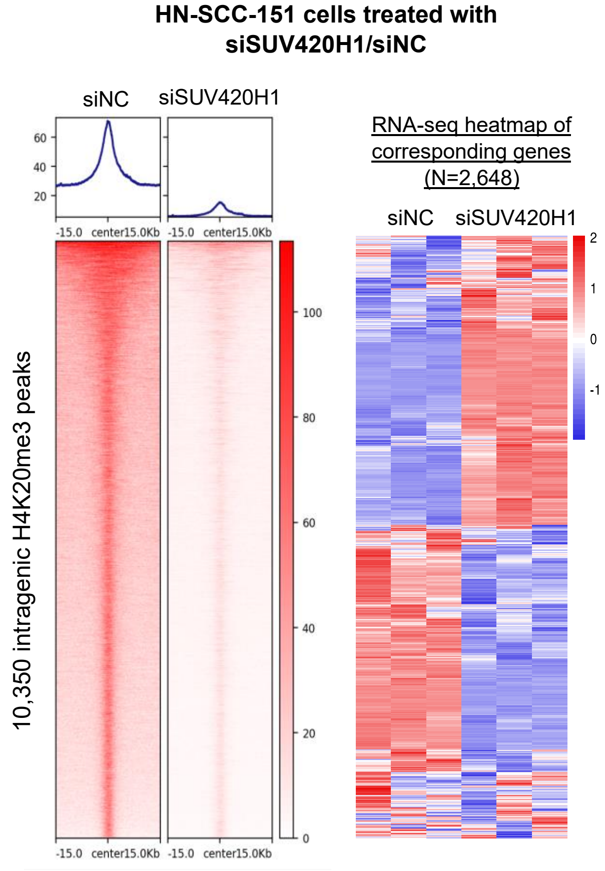

HN-SCC-151 cells treated with siSUV420H1/siNC
siNC
siSUV420H1
10,350 intragenic H4K20me3 peaks
-15.0
center
15.0Kb
RNA-seq heatmap of corresponding genes (N=2,648)
siNC
siSUV420H1
